# Functional Hierarchy of the Human Neocortex from Cradle to Grave

**DOI:** 10.1101/2024.06.14.599109

**Authors:** Hoyt Patrick Taylor, Khoi Minh Huynh, Kim-Han Thung, Guoye Lin, Wenjiao Lyu, Weili Lin, Sahar Ahmad, Pew-Thian Yap

**Affiliations:** Department of Computer Science, University of North Carolina, Chapel Hill, U.S.A.; Department of Radiology, University of North Carolina, Chapel Hill, U.S.A.; Biomedical Research Imaging Center, University of North Carolina, Chapel Hill, U.S.A.

**Author notes:** Corresponding authors: Hoyt Patrick Taylor IV and Pew-Thian Yap, **Correspondence** Correspondence and requests for materials should be addressed to P.-T. Yap.

## Abstract

Recent evidence indicates that the organization of the human neocortex is underpinned by smooth spatial gradients of functional connectivity (FC). These gradients provide crucial insight into the relationship between the brain’s topographic organization and the texture of human cognition. However, no studies to date have charted how intrinsic FC gradient architecture develops across the entire human lifespan. In this work, we model developmental trajectories of the three primary gradients of FC using a large, high-quality, and temporally-dense functional MRI dataset spanning from birth to 100 years of age. The gradient axes, denoted as sensorimotor-association (SA), visual-somatosensory (VS), and modulation-representation (MR), encode crucial hierarchical organizing principles of the brain in development and aging. By tracking their development throughout the human lifespan, we provide the first ever comprehensive low-dimensional normative reference of global FC hierarchical architecture. We observe significant age-related changes in global network features, with global markers of hierarchical organization increasing from birth to early adulthood and decreasing thereafter. During infancy and early childhood, FC organization is shaped by primary sensory processing, dense short-range connectivity, and immature association and control hierarchies. Functional differentiation of transmodal systems supported by long-range coupling drives a convergence toward adult-like FC organization during late childhood, while adolescence and early adulthood are marked by the expansion and refinement of SA and MR hierarchies. While gradient topographies remain stable during late adulthood and aging, we observe decreases in global gradient measures of FC differentiation and complexity from 30 to 100 years. Examining cortical microstructure gradients alongside our functional gradients, we observed that structure-function gradient coupling undergoes differential lifespan trajectories across multiple gradient axes.

---

Understanding how brain network organization changes across the human lifespan is a central and longstanding goal in neuroscience. Recent work has shown that functional connectivity is organized along axes manifesting as smoothly varying “gradients” on the cortex^1^. These gradients stratify cortical locations according to orthogonal connectivity patterns, capturing smooth variations in fundamental motifs of network architecture. These motifs reflect hierarchical features of brain networks, as evidenced by their alignment with established progressions in information processing^2–8^. These spatial progressions mirror well-defined hierarchical distinctions in sensory, associative, and integrative functions, supporting the interpretation of gradients as proxies for large-scale processing hierarchies.

Establishing a normative timeline of how these governing motifs of functional connectivity architecture develop in healthy individuals across the human lifespan is essential to a principled explanation of the onset, development, and timing of psychopathology^9^. Moreover, enumerating the lifespan trajectories of cortical regions with respect to global organizational motifs of the functional connectome provides rich insight into the topographic maturation of information processing hierarchies that scaffold complex cognition. By elucidating these developmental patterns, we can better comprehend how deviations from normative trajectories may contribute to neurodevelopmental and neurodegenerative disorders.

Recent work in this vein has challenged the conventional strategy of monolithically categorizing the function of discrete brain regions or networks, instead suggesting that network architecture is best understood in terms of the topographic organization of information processing motifs that smoothly vary as gradients across the cortex^2, 10^. While these motifs are referred to as functional hierarchies^2, 11, 12^, this terminology does not imply a strict causal or unidirectional flow of information. Rather, it reflects the observation that regions at the extremes of each gradient tend to play distinguishable roles—ranging, for example, from unimodal sensory processing to transmodal integrative functions. The most well-studied connectivity gradient is the canonical sensorimotor-association (SA) axis. Explaining the most variance in FC, the SA axis is anchored at one end by primary unimodal systems and at the other end by high-order association cortex coinciding with the default-mode network (DMN)^1, 2^. Similarly, there are repeated observations that the gradient explaining the second most variance in FC is an axis spanning between visual and somatosensory cortex (i.e., the VS axis), smoothly partitioning cortical locations according to their preferential involvement in modality-specific processing^1, 2^. Finally, many studies have identified a tertiary gradient of FC (i.e., the MR axis) anchored at opposite ends by cortical regions preferentially involved in modulation (frontoparietal control and attention networks) versus regions involved in representation (default, sensory, and visual networks)^1, 13, 14^.

Several studies have sought to characterize how one or more of these axes change during particular developmental periods, yielding observations that serve as a valuable foundation for establishing a multi-dimensional lifespan gradient atlas^15, 16^. Xia et al.^17^ demonstrated altered FC gradient ordering during childhood compared to healthy adults, with the VS axis explaining the most variance in FC, and an axis similar to the SA axis with important topographic differences constituting the secondary gradient of FC. Further, an age-based sliding window analysis of group-average FC gradients between 6 and 17 years found that convergence to the adult gradient ordering occurs during early adolescence, with higher-order association cortex undergoing more protracted gradient development than unimodal cortex^17^. Finally, one study analyzed the development of gradients during adulthood, finding that 3-dimensional gradient embeddings of canonical resting-state networks become increasingly dispersed during later adulthood, marking decreasing within-network coherence during aging^18^. While these studies have contributed a valuable foundation for understanding the development of FC gradient architecture, no research to date has offered a cohesive analysis of gradient architecture spanning from infancy to old age.

Concurrently, there is a vast body of non-gradient analyses of developmental changes in FC organization that provide invaluable context for gradient-based studies. Repeated observations demonstrate that, during infancy and early childhood, spatially dispersed, high-order resting-state networks are weakly expressed, with FC dominated by strong local connections in unimodal cortex^19–23^. During childhood, the basic substrates of complex cognition, including working memory, attention, self-referential thinking, and language ability, begin to emerge, necessitating the transition from a distinctly infant to an adult-like FC architecture. Graph-theoretic studies have established the importance of integration, segregation, and the modular architecture resulting from their interplay as being crucial to this transition, driving the emergence of higher-order systems including default and control networks, which serve as the principal facilitators of representation and top-down executive control, respectively^24–26^. Importantly, such transmodal networks globally coordinate information processing and are cortically distributed. As a result, their emergence and strengthening during childhood is enabled by increases in long-range connection density and strength, which mediate the transition from the highly local infant to global adult-like FC organization^27^ by the end of childhood and beginning of adolescence.

Once adult-like FC organization has been established, the honing and expansion of cognitive capacities incurred by adolescence and early adulthood coincide with subtle refinements of this architecture^28^. In particular, fine tuning of the integration and segregation in high order default-mode, frontoparietal control, and attention networks increase during this period, leading to more pronounced information processing hierarchies^29, 30^. During aging, degradation of the segregation-integration balance that scaffolds hierarchical FC architecture has been associated with cognitive decline^31–33^. Moreover, this period is characterized by global de-differentiation in FC that coincides with decreasing fluid intelligence and working memory.

Previous FC gradient studies provide useful insight into how gradient organization develops during particular developmental periods, however none of these studies encapsulate the entire lifespan and their analysis methods and reporting vary significantly. We aim to address these shortcomings of the developmental FC gradient literature by analyzing functional MRI (fMRI) data of healthy individuals spanning from neonates to centenarians in a consistent analytical framework. Under this framework, data across the lifespan is pre-processed identically, FC gradients are aligned to a common space that captures organizational axes of FC across the entire lifespan, and gradient value trajectories are modeled at the vertex level using generalized additive mixture models (GAMMs). We sought to reconcile, augment, and unify previous developmental FC studies by harnessing the power of the FC gradient framework in combination with a large sample size, high-quality fMRI dataset. We charted the lifespan trajectories of well-studied and validated axes of FC organization. We then used this lifespan chart of global FC architecture to characterize changes in the topography of individual gradient axes, global features, and multi-dimensional embedding the gradients collectively constitute.

## RESULTS

Our analysis aimed to chart changes in global intrinsic FC organization across the human lifespan. To this end, we computed low dimensional representations of the vertex-wise functional connectivity data of 3,556 subjects scanned at 3,972 unique time points with ages spanning from 16 days to 100 years using diffusion embedding^1^ (Figure 1). When applied to FC data, diffusion embedding has been demonstrated to yield smooth patterns of variation in connectivity, referred to as FC “gradients”. In tandem, diffusion embeddings preserve global connectivity relationships in the underlying FC graph, providing a low-dimensional approximation of the global organizing geometry of the functional connectome.

**Fig. 1.**
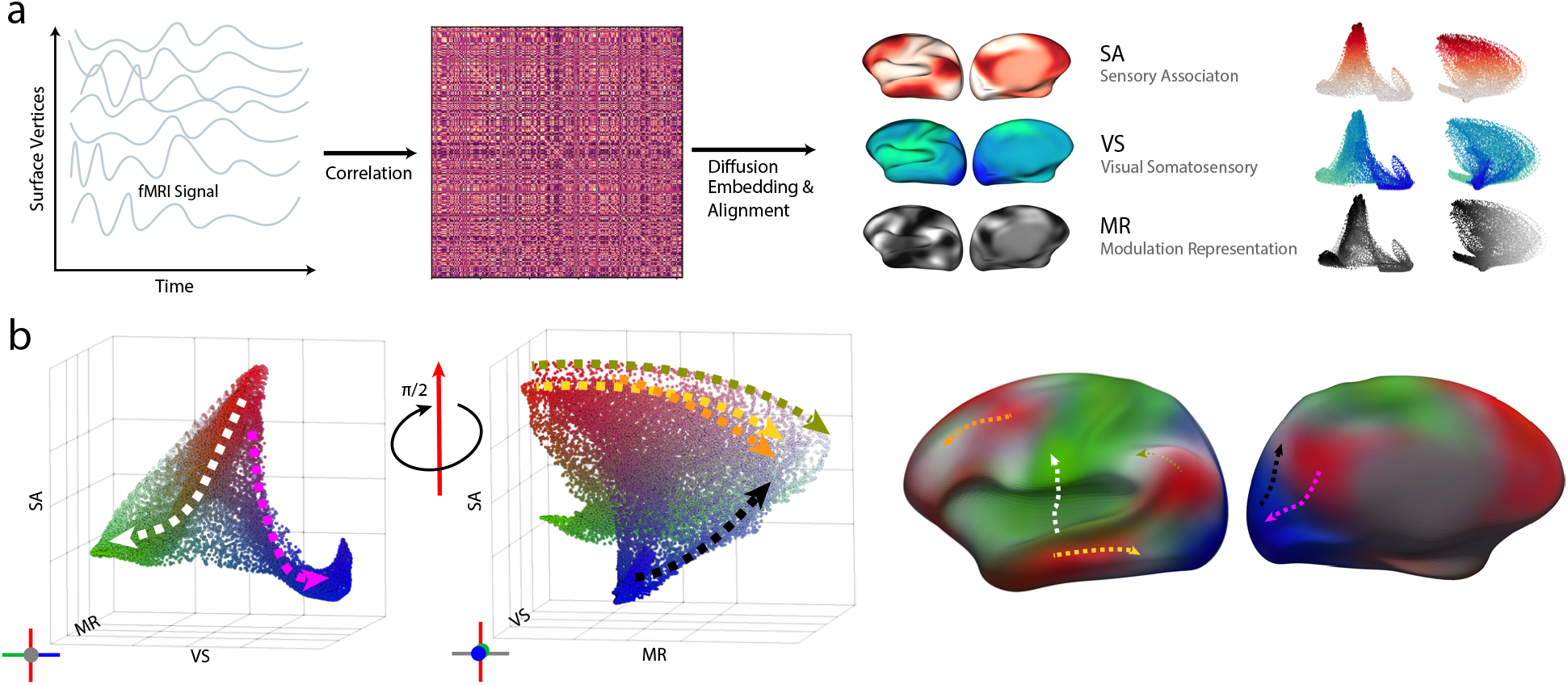
Overview of gradient manifold computation and its interpretation. **a**, For each subject, fMRI signal is mapped to the cortical surface and a functional connectivity (FC) matrix is computed via Pearson correlation coefficient. Individual subject FC gradients are obtained via diffusion embedding applied to the FC matrix, and are aligned to the template gradient axes of interest: sensory-association (SA), visual-somatosensory (VS), and modulation-representation (MR). Respectively, these axes differentiate cortical locations by their implication in association (red) versus unimodal (white), their preferential recruitment in either visual (blue) or somatomotor (green) domains, and their propensity to engage in top-down modulation (white) versus representation (black). **b**, The taxonomy of the adult gradient manifold in the embedding space (left) and the cortical surface (right) with a unified color map that combines the three used in **a**. Several paths along the gradient manifold are displayed with corresponding color-coded paths along the cortical surface, demonstrating the cortical realization of hierarchies enumerated by the gradient manifold.

We employed a framework that allows for joint analysis of the FC gradients for all subjects across 5 datasets with ages spanning the entire lifespan in a common space. First, we computed a template gradient set by applying weighted principal component analysis to the set of all subject gradients. These template gradients corresponded to well-studied cortical gradients spanning respectively between cortical locations implicated in sensorimotor-association (SA), visual-somatosensory (VS), and modulation-representation (MR). Together, these axes define the embedding space in which we carried out the remainder of our analysis. We proceeded by aligning all individual subject gradients to this template space using Procrustes alignment^34^, after which the gradient value at each vertex for all subjects measures the same connectivity characteristic. Finally, by harmonizing gradient values across the lifespan to control for cohort effects and reduce heteroscedasticity and fitting generalized additive mixed models (GAMMs)^35^ to gradient value at each cortical surface vertex versus age, we obtain a population-level estimate of the trajectory of these aligned gradient values across the entire lifespan.

Our lifespan gradient fit provides the first ever comprehensive dynamic atlas of global FC organization from days after birth to 100 years. Age-related changes in the cortical topography of each gradient provide insight into the spatial realization of three crucial organizational hierarchies on the human cortex. Simultaneously, the 3 dimensional gradient embedding reveals how the interrelationships between these hierarchies develop from birth to death. More specifically, the position of a vertex in the lifespan gradient embedding at a given age encodes its position with respect to the three principal axes of FC organization, giving a detailed description of that vertex’s connectivity characteristics. Our result provides a temporally dense and spatially detailed normative reference of how these crucial information processing motifs evolve with respect to cortical locations.

### Changes in gradient topography are concentrated in infancy and early childhood

A central objective of this study was to characterize how the cortical topography of the sensorimotor-association (SA), visual-somatosensory (VS), and modulation-representation (MR) axes vary with age. To this end, we analyzed our vertex-wise GAMMs of gradient values mapped to the cortical surface (Figure 2a). This allows us to examine the smooth temporal development of the principal gradients of FC across the human lifespan. The cortical topography of each gradient provides crucial insight into the spatial realization of critical patterns of variation in connectivity. The distribution of values for each gradient, depicted via a density plot, details the hierarchical architecture of its corresponding connectivity characteristic and allows for the characterization of global changes in each gradient. Peaks in gradient value density plots arise when many vertices share the same FC profile with respect to a gradient, whereas uniform, widespread density distributions indicate highly heterogeneous or stratified FC. Notably, gradient topography changes most significantly during the first 4 years of life, consistent with observations that infancy and early childhood represent critical periods for the development of large-scale adult-like network architecture^21, 23, 36–38^.

**Fig. 2.**
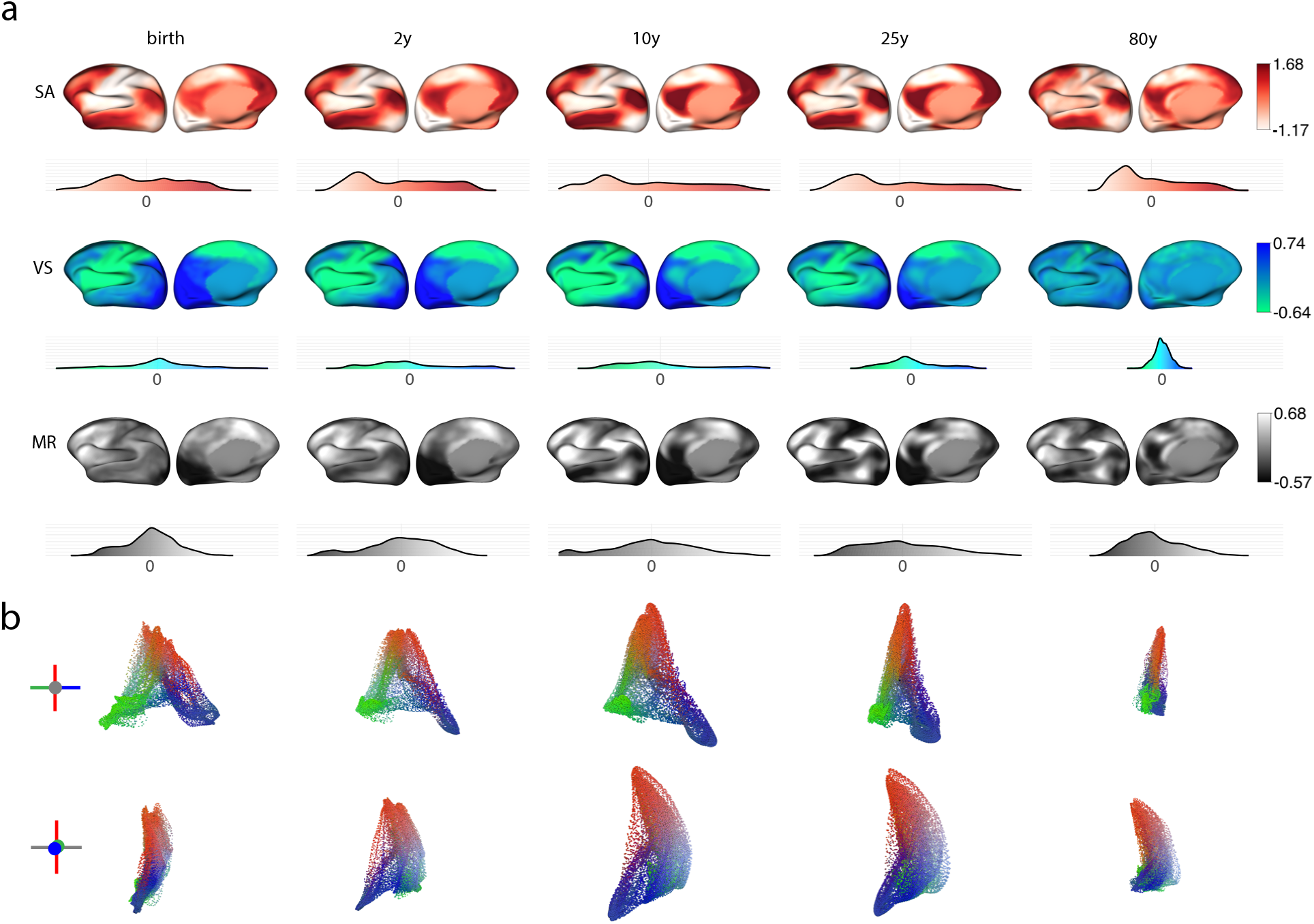
Gradient organization across the human lifespan. Sensory-association (SA), visual-somatosensory (VS), and modulation-representation (MR) gradients of FC undergo differential development across the human lifespan. **a**, GAMM fits of FC gradient value are plotted on the cortical surface with density plots displaying gradient value distribution at selected timepoints. **b**, Embedding plots given by gradient GAMM fits in SA-VS plane (top) and SA-MR plane (bottom). Timepoints were selected to reflect key developmental milestones across the lifespan.

We also quantitatively assessed the significance of age-related change in gradient value across the cortex on a region-wise basis by fitting GAMMs (*k* = 6 basis functions for smooth age term) to the mean harmonized gradient value within each of the 400 regions of the Schaefer parcelletion^39^. SA gradient value varied significantly across the lifespan in the majority of cortical regions (*p*_FDR_ *<* 0.05 for 99.5% of regions), with the strongest age-related changes concentrated in somatosensory and default-mode parietal cortices. VS gradient value also varied significantly for a slightly greater proportion of cortical regions (*p*_FDR_ *<* 0.05 for 100% of regions), with fits of highest significance occurring in visual and somatosensory cortex. Age-related changes in MR gradient value were similarly significant across cortical regions (*p*_FDR_ *<* 0.05 for 97.25% of regions), with maximal significance occurring in control-oriented parietal and frontal cortices. We observed significant non-linearity in the lifespan trajectory of SA, VS, and MR gradient value across most cortical regions, with maximal lifespan fluctuation taking place at vertices occupying gradient poles.

The mature SA axis describes the principal organizing motif of functional connectivity, stratifying cortical locations according to their position along a hierarchy spanning from primary sensory regions to heteromodal association cortex^1^. In adults, we observe homogeneous values at the unimodal end of this axis and highly stratified values on the association association pole (Figure 2a, SA density plot for 25 years). This is further evidenced by smoothly decreasing SA values emanating from the highly localized association focal points (dark red) and homogeneous sensory values (white) in unimodal cortices (Figure 2a, SA surface plots for 25 years). Between mid-adolescence and old age, the cortical topography and heavy-tailed distribution of the SA axis remain consistent, indicating that the spatial layout of this hierarchy is established during infancy, childhood, and early adolescence.

During early life, SA organization is distinct from that of adulthood, with very little stratification among cortical regions on its association pole and relatively greater stratification on its unimodal pole. The first two timepoints (birth and 2 years) in Figure 2a illustrate this difference, with homogenous values at the association pole (red) on the cortical surface, steep falloff in SA value density on the association side, and more gradual falloff in value density on the unimodal side. These distinctive features of the SA axis during early life underscore the absence of well-ordered sensory-association hierarchy and further indicate that FC organization during this period is dominated by unimodal processing and short-range connectivity, as regions at either pole of this ‘proto-SA’ axis tend to be spatially contiguous. Figure S1 illustrates the immature and distinct topography of the SA axis during infancy, while Figure S2 displays the convergence of SA topography to its mature form that plays out between 2 and 6 years.

The transition from a highly local and unimodally driven architecture to the stratified hierarchical architecture of the mature SA axis takes place between 2 and 10 years of age (Figure 2a, SA axis). During this period, we observe significant focal tuning of the association pole reflected by an increasingly heavy-tailed SA value distribution (Figure S10). This change is realized on the cortical surface as increasingly localized association (red) peaks that converge towards the canonical loci of the default mode network. These shifts coincide with the development and crystallization of crucial aspects of identity, cognitive ability, and personality that take place during this critical period^21, 28, 30, 40, 41^.

Like the SA axis, the mature modulation-representation (MR) axis enumerates an information processing hierarchy whose apex is comprised by spatially distributed heteromodal cortical regions. Whereas regions atop the SA axis are central to the DMN, the modulation pole of the MR axis is comprised of core constituents of the frontoparietal control network (FPC) and attention networks^14, 42^. Like the SA axis, the modulation pole of the MR axis is highly stratified (Figure 2a, 25 years, MR, white). A peak in the center of the MR distribution in combination with gradually decaying tails at both poles indicate that only a small number of brain regions are recruited strictly for modulation or for representation. Rather, the plurality of cortical regions occupy some middle ground between these extremes. As a result, the spatially dispersed modulation and representation poles of the MR axis are highly localized.

During early life, the MR axis very weakly resembles its mature form (Figure 2a, MR, birth + 2 years). Figure S1 and Figure S2 illustrate the maturation of the MR axis during early life, exhibiting the least similarity to its mature form among our three gradients of interest. Neither modulation or representation poles are highly localized, with relatively low degrees of stratification towards either extreme. Between birth and 2 years, superior parietal regions encompassing portions of the mature dorsal attention network as well as inferior parietal lobule constituting loci of the salience and frontoparietal control networks move towards the modulation pole, however there is still weak stratification at 2 years. Between 2 and 10 years, adult-like MR topography emerges with increasing stratification of both poles (Figure S2), indicating that this period is critical to the development of the differentiation between control-oriented and representation-oriented cortical regions. Interestingly, the visual cortex is established as the representation pole of the MR axis during the first year of life, with the somatosensory cortex migrating to the representation pole much later during early adulthood. This distinction suggests that representation-oriented coordination between visual and somatosensory cortex coincides with the development of mature, adult-like network architecture seen during adolescence. The MR axis undergoes further refinement between 10 and 25 years, with the somatosensory cortex moving towards the representation pole, and peaks in the frontal cortex shifting towards the modulation pole (Figure S3). This prolonged developmental trajectory is consistent with repeated observations that development of executive function and control is protracted and persists into early adulthood^43, 44^.

In contrast to the high-order information processing hierarchies corresponding to the SA and MR axes and the complex topographic development they undergo, the VS axis differentiates brain regions with respect to preferential involvement in the two primary unimodal processing domains and has largely stable cortical topography throughout the lifespan. The range of the VS axis is maximal during early life (peaking at 6.8 years), indicating that the distinction between unimodal sensory processing modalities is dominant in shaping FC organization during this period. At birth, the distribution of regions along the VS axis is symmetrical (Figure S1). During the first two years of life, values become increasingly dense on the somatosensory pole of the VS axis, indicating increased integration among motor regions. Simultaneously, stratification of values on the visual pole of the VS axis increases from birth until 10 years, indicating that functional differentiation of the visual processing system is ongoing throughout this period. The most prominent topographic change in the VS axis across the lifespan is in the migration of dorsal attention regions in the superior parietal cortex towards the visual pole and of salience and of auditory regions towards the somatomotor pole. These changes take place gradually between birth and 10 years, with more rapid migration of regions in auditory and salience networks towards the somatomotor pole of the VS axis taking place during later childhood. This topography is suggestive of differential roles of the dorsal and salience/ventral attention networks with respect to visual and somatomotor processing. The global range and the stratification of extreme visual and somatosensory extreme values of the VS axis decrease significantly and monotonically across the lifespan after 6.8 years old, signifying that the distinction between visual and somatosensory processing streams becomes a less formative feature of global network architecture during adolescence and adulthood. These changes, when considered alongside the topographic changes exhibited by the VS axis, indicate a convergence toward a more domain-agnostic and multi-modally integrated connectivity architecture that plays out between birth and cognitive maturity.

### Global measures of FC gradient organization exhibit non-linear changes across the lifespan

We observe significant non-linear developmental trajectories in global properties of all three of our gradient axes and the 3-dimensional embedding they constitute. To assess the global expression of a gradient, we calculate its range using the inter-vigintile spread (5th to 95th percentile). This provides a robust measure of the prominence of the corresponding connectivity motif. We fit GAMMs to the ranges of the SA, VS, and MR axes and observe significant non-linear gradient-specific developmental trajectories across the lifespan, indicating complex FC reorganization in multiple hierarchical dimensions (Figure 3). Both SA and MR axes undergo gradual expansion throughout infancy, childhood, and adolescence. The scale of the SA axis reaches a pronounced peak during early adulthood at 24.9 years (95% CI 23.0–26.8 years). Similarly, the range of the MR axis increases monotonically throughout infancy, childhood, and adolescence, reaching a more protracted and smooth peak at 24.3 years (95% CI 22.3–26.3 years). During early adulthood, the range of the SA and MR axes are stable, however the SA axis undergoes more rapid contraction during mid and late adulthood compared to the MR axis, which only contracts modestly. Both SA and MR axes exhibit inverted U-shaped development previously observed in lifespan FC development^45^. The VS axis, which differentiates between the primary unimodal processing centers, obtains its maximum range during late childhood at 6.8 years (95% CI 6.5–7.0 years), undergoing gradual contraction throughout the remainder of the lifespan.

**Fig. 3.**
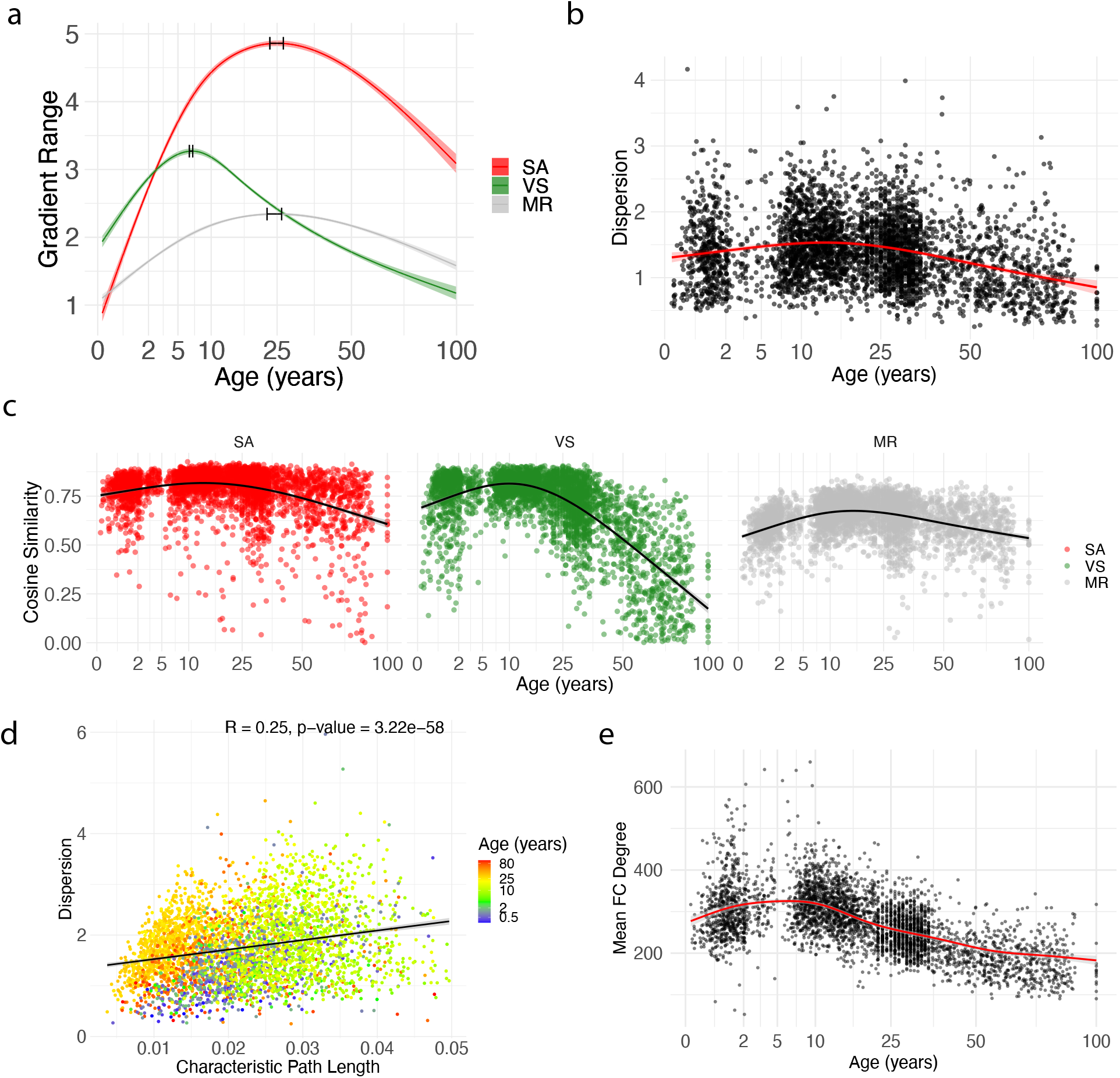
Global gradient metrics denote changes in hierarchical FC architecture across the lifespan. **a**, Inter-vigintile gradient range versus age (log scale) with GAMM fits for SA, VS, and MR axes. **b**, Gradient dispersion versus age with GAMM fit. **c**, Cosine similarity between individual subject gradients and template gradient set used for alignment. **d**, Characteristic path length versus gradient dispersion. **e**, Mean functional connnectivity degree across the lifespan with GAMM fit. Shaded areas indicate the standard error for all fits.

Expansion of the transmodal SA and MR gradients throughout development is consistent with abundant evidence that heteromodal association and modulatory attention and executive cortex have protracted maturation^43, 46–48^. As gradient range measures the differentiation in FC between cortical locations at each pole of the gradient, the expansion of the SA axis signifies increasing functional dissimilarity between the default-mode association cortex (Figure 1a, top right cortical surface, red) and the primary unimodal cortex (Figure 1a middle right cortical surface, blue and green). In turn, the expansion of the MR axis marks an increasing contrast in the connectivity profiles of modulation-implicated (frontoparietal control and attention) cortices and representation-implicated (default, sensory, visual, and limbic) systems. The VS gradient displays highly divergent global development compared to the SA and MR axes. Namely, the range of the VS axis peaks during childhood at 6.8 years, followed by significant decline thereafter. Figure 2 gives two alternate depictions of the VS axis contraction as given by the shrinking tails of the gradient value density distributions (Figure 2a) and the horizontally contracting embedding plots in the SA-VS plane (Figure 2b, top).This trend is consistent with previous studies that observe a decrease in the variance explained by the VS axis^38^ as well as studies showing a transition from local to long-range FC organization throughout development^27^.

We also studied how the dispersion of vertices in our 3-dimensional embedding space changed throughout the lifespan by computing gradient dispersion, the average embedding distance between each vertex and the embedding centroid for each subject. Because Euclidean distance in the diffusion embedding space approximates geodesic distance in the underlying FC matrix, gradient dispersion provides a measure of global FC differentiation with respect to the SA, VS, and MR axes. High dispersion values indicate globally heterogeneous FC profiles, while lower dispersion values correspond to lower cortex-wide diversity in FC profiles. It is important to note that dispersion provides a scalar summary of the spread of vertices in all three gradient dimensions despite the differential contribution of each axis to the Euclidean distances in embedding space. With this in mind, dispersion should be interpreted in combination with information about the relative scale of each gradient axis (Figure 3a).

We fit a GAMM to model the trajectory of mean dispersion with respect to age (Figure 3). Age was treated as a smooth term modeled using smooth splines with *k* = 5 basis functions. Figure 3b shows a steady increase in dispersion during infancy and childhood, with rapid increases in the SA and MR axis ranges and modest decline in the VS axis range (Figure 3a). Dispersion reaches its lifespan peak at 13.8 years (95% CI 11.3–16.3 years), remains relatively stable through early adulthood (25 years), and decreases steadily thereafter. Increases in dispersion throughout infancy, childhood, and early adolescence coincide with significant expansion of the SA and MR axes and moderate contraction of the VS axis, reflecting a global increase in FC differentiation, particularly with respect to high-order information processing hierarchies. The peak in dispersion occurs prior to the peak range in SA and VS axes due to an increasing reduction in the range of the VS axis which begins at roughly 12 years old. Declining dispersion during later adulthood reflects decreasing global connectivity differentiation in tandem with reduction in the ranges of all three gradient axes, indicating that FC profiles become more globally homogeneous during aging. This finding is consistent with prior studies that observe decreasing functional segregation during aging^49, 50^. This nonlinear trajectory serves as a normative reference for the global differentiation of FC with respect to the first three hierarchical axes of organization across the lifespan.

To further substantiate our gradient-based global measures of FC organization, we computed a number of graph-theoretical metrics from the FC matrices underlying our gradients and examined their lifespan trajectories and relationships to gradient metrics. Notably, characteristic path length, an indicator of the average shortest path between nodes, exhibited moderate but highly significant correlation with gradient dispersion (Figure 3d). Because lower characteristic path length is associated with higher global efficiency ^51^, this finding suggests that gradient dispersion captures aspects of network integration and efficiency in the developing functional connectome. Consistent with prior lifespan studies^52, 53^, we observed that young adults had lower characteristic path length than adolescents, while infants, children, and older adults showed intermediate values. We also computed the spatial mean of FC degree (i.e., vertex-wise sum of thresholded FC matrices used as input for diffusion map embedding), which increased steadily from infancy to a peak at approximately 10 years before declining. Interestingly, this developmental trend did not parallel the range of any specific gradient axis, implying that gradient metrics capture more nuanced network characteristics beyond simple FC strength.

To compare FC gradients across subjects, we aligned individual gradient sets to a template gradient set. This template was derived using PCA on the gradients of all subjects and included the SA, VS, and MR axes. To minimize age-related bias, we applied a weighted PCA scheme when computing the template gradients. While these three axes capture the most prominent FC motifs across the lifespan, individual gradients vary significantly in their topography, ordering, and scale. To probe how the topography of individual gradients varied with respect to our canonical template gradients, we computed the cosine similarity between the aligned gradients of each individual and the corresponding template gradient. Figure 3c displays template cosine similarity for all subjects versus age with GAMM fits analogous to dispersion. The similarity of each axis to the lifespan template increased significantly between birth and early adolescence, with maximum similarity to the template achieved for the VS axis at 9.8 years (95% CI 9.5–12.3 years), for the SA axis at 13.5 years (95% CI 12.3–14.8 years), and finally for the MR axis at 15.8 years (95% CI 14.8–16.8 years). This ordering reflects the expected maturational timing of our gradient axes, with the most simplistic and concrete (VS) axis maturing first and the axis most implicated in high-order executive function (MR) maturing last, aligning well with prior studies on the development of adult-like network architecture at the onset of adolescence^50^. The decline in cosine similarity to our template observed in all three axes that occurs after mid adolescence reflects increasing diversity in global functional connectivity architecture. Further, the sharp decline in VS axis similarity may indicate that FC organization becomes less organized around sensory modality during aging.

### Resting-state networks undergo significant gradient reorganization throughout the lifespan

Previous studies have demonstrated that canonical resting-state networks are stratified along the primary FC gradients^1, 54^. In tandem, there is mounting evidence that brain development plays out along cortical gradients including the SA axis^46, 50^, with unimodal networks achieving maturity far earlier than limbic, attention, and association networks^40, 55, 56^. We investigated the reconfiguration of canonical resting-state networks in our aligned gradient embedding space using Yeo et al.’s FC-based 7-network parcellation^57^. We first examined network trajectories in the embedding space by computing the mean gradient value for each network and gradient across subjects, reflecting the positioning of network-specific vertices along each gradient axis. To evaluate the lifespan trajectory of networks along each gradient axis, we fitted separate GAMMs to the gradient means of each network. Assessing based on gradient means, the networks were intuitively ordered with respect to each axis, with the default and fronto-parietal control networks occupying the association and modulation extremes of the SA and MR axes, respectively, and the visual and somatosensory networks occupying their respective poles of the VS axis. Importantly, we observe differential trajectories of gradient means across networks and across axes (Figure 4a), indicating that networks undergo complex maturational timing with respect to the principal axes of FC organization.

**Fig. 4.**
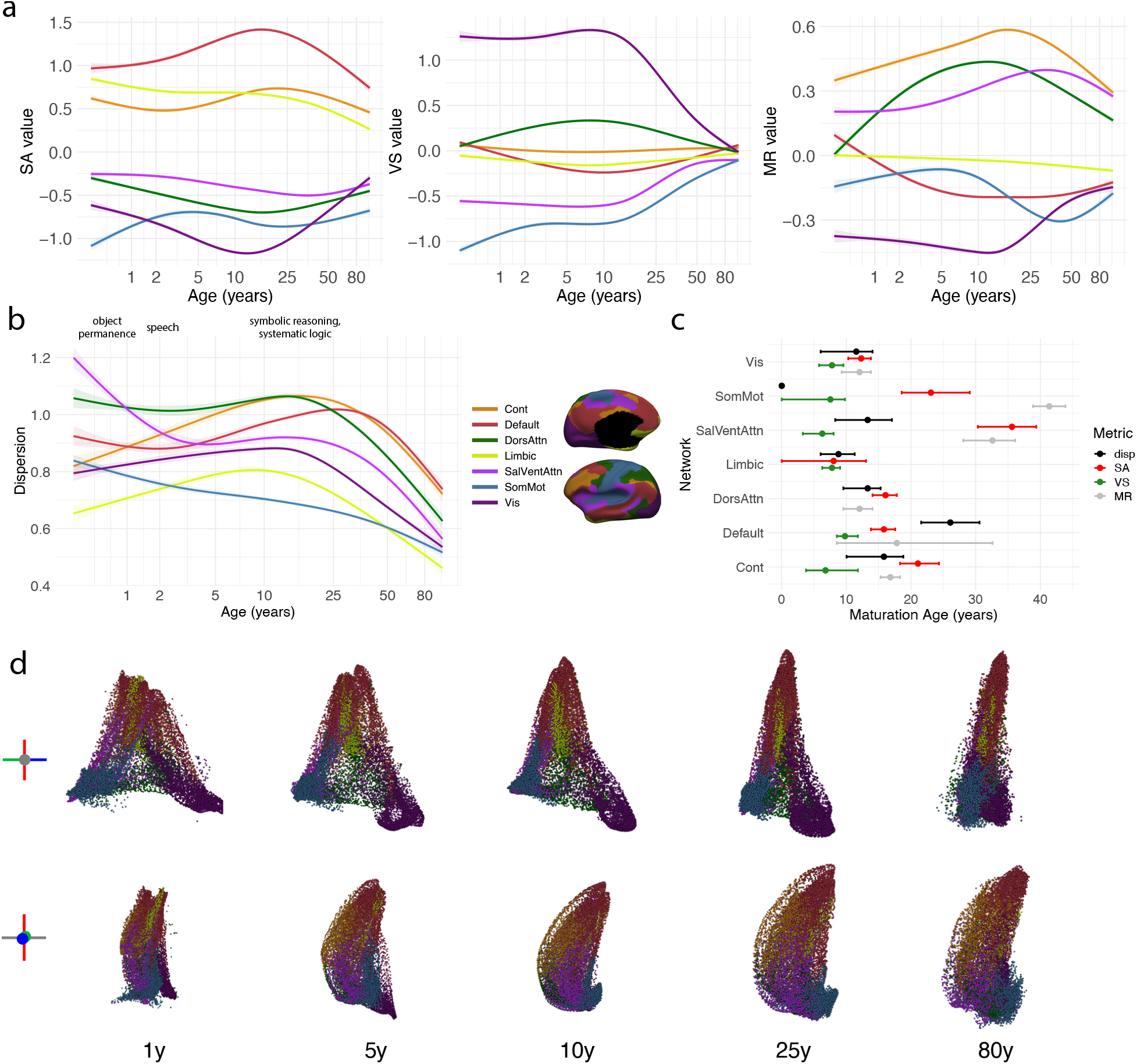
Lifespan development of functional gradients with respect to canonical resting-state networks across the lifespan. **a**, GAMM fits of mean gradient value versus age within each network parcel for SA, VS, and MR axes. **b**, GAMM fit of within-network dispersion versus age, computed as the mean squared Euclidean distance between each vertex’s coordinate in the SA-VS-MR gradient embedding space and the network centroid. **c**, Maturation age of network-wise gradient value and dispersion. For gradient value, maturation age is taken as the timepoint at which the GAMM fit for a particular network along a particular gradient reaches its most extreme value along that gradient. For dispersion, maturation age is treated as the age at which maximum dispersion occurs for a network. **d**, GAMM-fitted gradient embedding plots colored using the Yeo 7-network parcellation at selected timepoints across the lifespan viewed from the SA-VS plane (top) and the SA-MR plane (bottom).

The expansion of the SA axis during development and its contraction during aging mediate the lifespan changes in resting-state network configuration along this unimodal-to-transmodal axis. Namely, default and control networks move towards the association pole of the SA axis between birth and mid-adolescence, indicating that the establishment of the SA axis coincides with reorganization of these high order networks (Figure 4a, left). Conversely, the salience/ventral attention, dorsal attention, and visual networks move down the SA axis gradually between birth and 25 years, and move to a more central position thereafter. Migration towards the sensory pole of the SA axis indicates that these networks become more closely integrated with the unimodal “base” of the sensory-association information processing hierarchy. Interestingly, the somatomotor network begins at the extreme sensory end of the SA axis, moves to a more central (less unimodal) position between birth and 3 years, and then converges to the trajectory observed in the attention and visual networks (Figure 4c). This differential trajectory of the somatomotor network during early development is potentially explained by consideration of the topographic reorganization of the SA axis during this period, wherein the sensory pole is initially strongly concentrated in the somatomotor cortex and gradually equalizes with the visual system thereafter (Figure 2a).

The VS axis discriminates brain locations according to preferential involvement in visual versus somatosensory processing streams. Thus, heteromodal networks including default, control, limbic, and dorsal attention occupy central positions on the VS axis throughout the lifespan (Figure 4c). Visual and somatomotor networks occupy the opposite extreme poles of the VS axis throughout the lifespan, with the salience/ventral attention network situated on the somatomotor side of the axis. This positioning of the salience/ventral attention network is striking, indicating dominance of somatomotor processing in top-down attention modulation. During infancy, the somatomotor and visual networks are maximally distant on the VS axis (Figure 4a, middle). Notably, the mean VS coordinate of somatomotor vertices takes on its most extreme position at birth and trends towards a more central position throughout the duration of the lifespan. In contrast, the visual network occupies the extreme visual pole of the network at birth and throughout development, indicating that the visual network originates as highly differentiated along this axis. The salience/ventral attention network, which contains crucial areas of premotor cortex, also moves towards the somatomotor pole of the VS axis between birth and 10 years, indicating an increase in preferential involvement in somatomotor unimodal processing during this period. The networks occupying the poles of the VS axis (visual, somatomotor, and salience/ventral attention networks) are maximally separated along the axis between birth and 10 years and contract towards a central position rapidly thereafter in tandem with the global contraction of the VS axis as indicated by its declining gradient range (Figure 3a).

Network centroids along the MR axis exhibit the most complex lifespan dynamics, with varied maturational timings and developmental profiles across multiple networks (Figure 4c). The MR axis differentiates brain regions with respect to their preferential involvement in topdown modulation versus representation. It therefor stands to reason that control and attention networks occupy the modulation pole with default,and unimodal networks occupying the representation pole. Default and control networks become increasingly separated on the MR axis between birth and early adulthood. However, the default mode network reaches its mature extreme position on the modulation pole during early adolescence whereas the control network moves towards the modulation pole throughout adolescence. During adulthood, both default and control networks move to more central positions, coinciding with the contraction of the MR axis during this period (Figure 2b). Dorsal and ventral attention networks undergo differential development along the MR axis. The dorsal attention network reaches its maximum positions on the modulation pole during early adolescence, while the salience/ventral attention network undergoes more protracted development, reaching its maximum value during mid adulthood.

### Interactions between resting-state networks in gradient space track developmental refinement of FC organization

Our analysis of changes in dispersion and gradient value within restingstate networks demonstrated that gradient reorganization is nonlinear across the lifespan and heterogeneous across networks. Further, inspection of our GAMM fits of gradient value revealed that the 3-dimensional distributions of network nodes in embedding space change significantly across the lifespan. To characterize these changes and the interaction between networks in greater detail, we analyzed the pairwise distance distributions between vertices in each pair of networks. Previous studies have computed “between-network dispersion” by first taking the mean embedding coordinates of vertices within each network and then computing the Euclidean distance between these network centroids^18^. While this is a good first-order approximation, it discards rich information about the shape and spread of vertices within each network. Instead, we define the between-network dispersion as the mean distance between all pairwise Euclidean distances between vertices in two networks and normalize this quantity by the global dispersion of the embedding. We computed between-network dispersion for each network pair in each subject’s aligned gradient embedding, then fitted a GAMM versus age for each network pair with age (Figure 5).

**Fig. 5.**
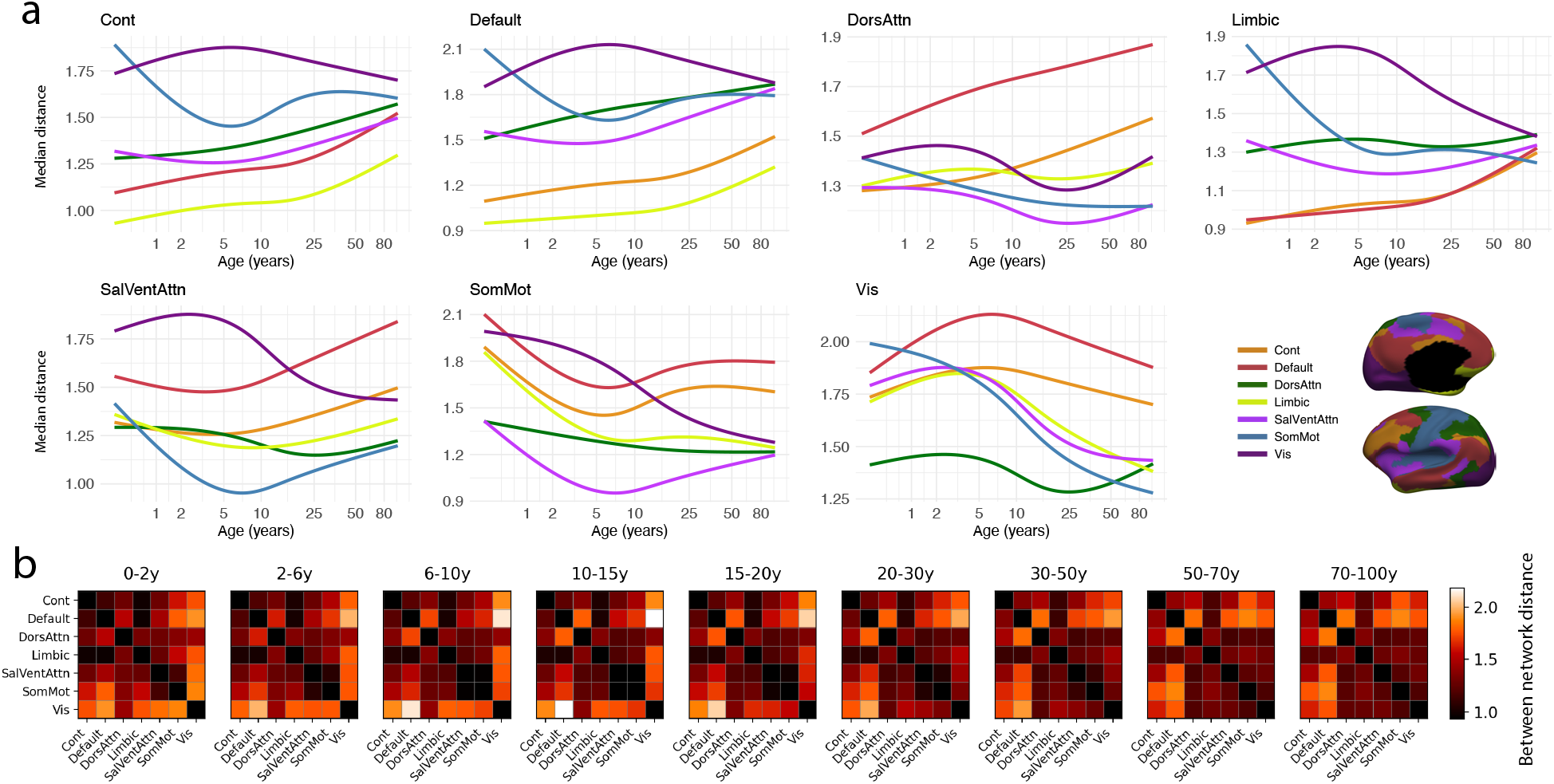
GAMM fits of mean gradient embedding distance between resting-state networks. Larger distances correspond to higher levels of functional differentiation and segregation. Small distances indicate functional similarity and integration. **a**, each plot displays the lifespan GAMM fit of the mean Euclidean distance in the SA-VS-MR gradient embedding space between vertices in the title network and vertices in each other network. **b**, Matrix plots of average network-network mean embedding distance in selected temporal windows, with lighter values denoting larger distance.

Our relative between-network dispersion metric can be interpreted as a measure of between-network segregation, with higher values indicating segregation between two networks and lower values indicating increased integration. With this in mind, Figure 5 illustrates the lifespan changes in integration/segregation for each pair of resting-state networks. We observe a period of rapid changes in between network dispersion between birth and 5 years, with increasing dispersion between visual-default, visual-control, and visual-dorsal attention and decreasing dispersion between somatomotor-default and somatomotor-control. These rapid reconfigurations during early life are consistent with evidence that infancy represents a critical period for the simultaneous proliferation and pruning of functional connections^58, 59^. Dispersion between the default and dorsal attention network increases throughout the lifespan, whereas the default and salience/ventral attention network undergo persistent differentiation only after 5 years. Simultaneously, dorsal attention and control networks become increasingly dispersed throughout the lifespan, indicating progressive segregation between these systems which is persistent throughout development.

The dispersion between default and control networks increases monotonically throughout development, with more rapid differentiation occurring after 10 years. This result indicates that the onset of differentiation between these networks is delayed until late childhood. Interestingly, somatomotor-control and somatomotor-default segregation decrease during early development, suggesting that the establishment of gradient topography during this period occurs in tandem with increasing coordination between high-order networks and the somatomotor network. This is potentially attributable to the importance of the acquisition of gross and fine motor skills during this period.

### Structure-function coupling undergoes nonlinear and divergent lifespan development across multiple cortical hierarchies

We also sought to probe the relationships between cortical microstructure and our FC gradients across the human lifespan. To this end, we computed structural gradients from affinity matrices based on pairwise Pearson correlation of cortical features, including cortical thickness, myelination, volume fractions (intra-cellular, extra-cellular, and intra-soma), and diffusion metrics (fractional anisotropy, mean diffusivity, microscopic fractional anisotropy, microscopic mean diffusivity, microscopic anisotropy index, and orientation coherence index). Structural gradients were then computed for each subject using the same approach as functional gradients, yielding a set of structural organization axes across all subjects. We then aligned the structural gradient embeddings of each subject to their corresponding functional gradients, which were previously aligned to our lifespan functional gradient template consisting of the SA, VS, and MR axes. We observed modest topographical similarity between structural gradients and individual SA, VS, and MR functional gradients, with an average spatial correlation between structural and functional gradient counterparts of *ρ* = 0.45 for the SA axis, *ρ* = 0.41 for the VS axis, and *ρ* = 0.38 for the MR axis. These modest spatial correlations suggests that the principal axes of functional organization are only moderately rooted in the underlying cortical microstructure. To assess lifespan changes in the similarity between our structural and functional gradients, we computed cosine similarity between aligned structural and functional gradients for each subject and subsequently fitted GAMMs to obtain lifespan trajectories of axis-specific structure-function coupling (Figure 6b).

**Fig. 6.**
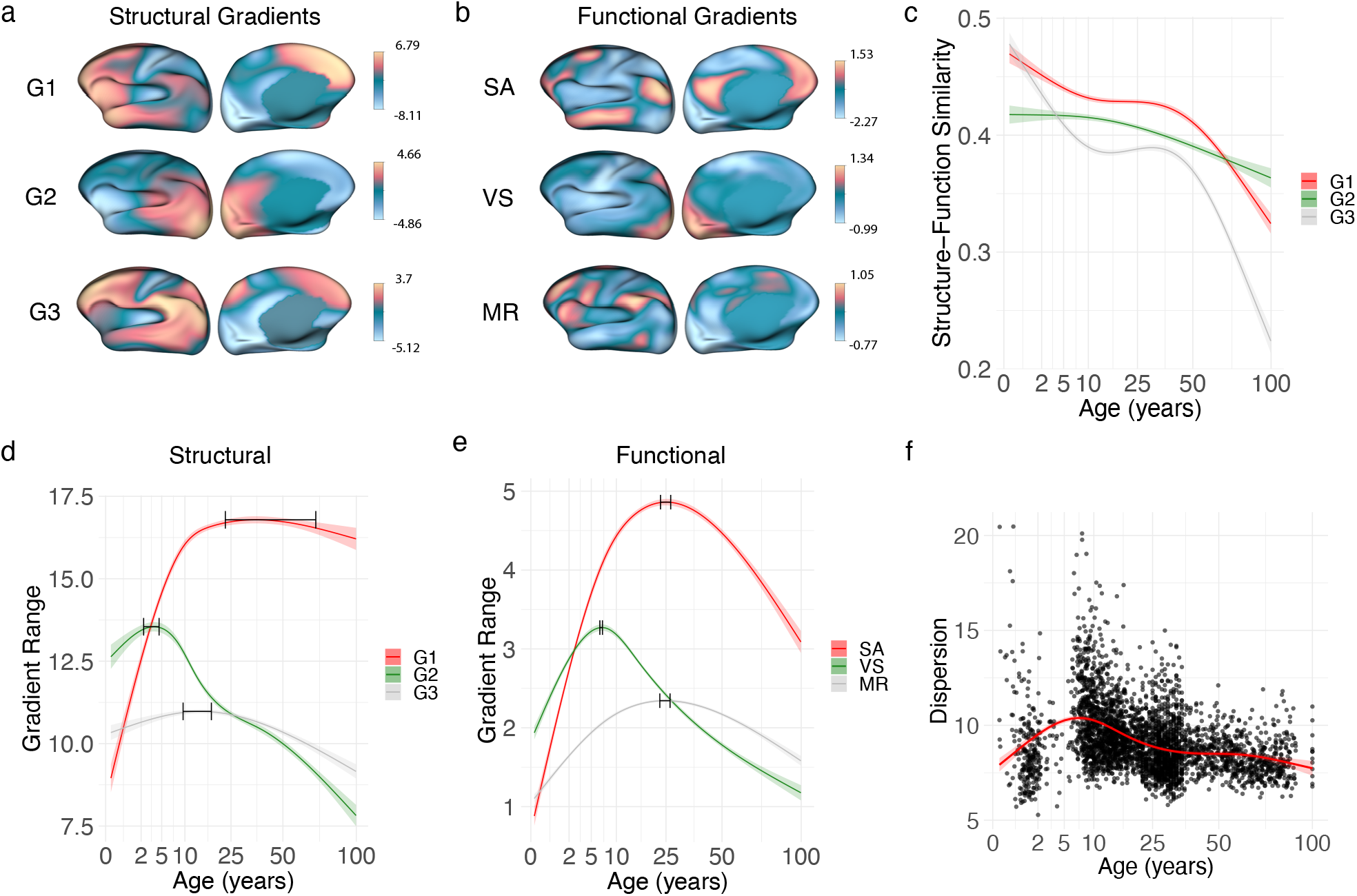
Analysis of structural gradients based on cortical thickness, myelination, and microstructural indices. **a**, Surface plots of group-level structural gradients. **b**, Functional gradient template (right). **c**, GAMM fits of structure-function coupling between individual structural gradients and individual functional gradients after Procrustes alignement for SA, VS, and MR axes. **d**, GAMM fits of gradient ranges for structural gradients. **e**, GAMM fits for gradient ranges of functional gradients. **f**, Dispersion of structural gradients from birth to death. The 95% CI for the peak of each trajectory is shown for **d** and **e**. Shaded areas indicate the standard error for all fits.

Not surprisingly, our structural gradients are topographically distinct from our functional gradients even after Procrustes alignment (Figure 6a). Qualitatively, structural and functional counterparts of the SA axis are similar, while the VS-aligned structural gradient exhibits large differences from the functional VS axis. Specifically, while the visual cortex is at the visual extreme of both the functional and structural VS axes, the somatosensory cortex does not reside at the somatomotor pole in the VS-aligned structural gradient. This indicates that differences in cortical microstructure between the visual and somatosensory systems do not topographically coincide with the functional VS axis. The MR-aligned structural gradient also diverges from its functional counterpart, with far less spatial localization in its modulation pole. It further lacks the representation pole in DMN loci in the inferior parietal lobule exhibited in the functional MR axis. Together, these results suggest that there is strong microstructural underpinning of the SA axis, but far weaker orthogonal gradients of cortical microstructure underpinning the secondary and tertiary VS and MR axes.

Developmentally, the topographic similarity between these structural gradients and their functional counterparts undergo significant nonlinear change across the lifespan, with structurefunction coupling decreasing with age (Figure 6c). Structure-function gradient similarity decreases rapidly for the SA and MR axes during infancy and early childhood, indicating that the functional differentiation underpinning the development of these functional hierarchies is not accompanied by the development of equivalent microstructural gradients. Strucure-function coupling for the VS axis is relatively stable throughout infancy and childhood, exhibiting modest decline after 10 years. Lifespan trends in structural gradient range (Figure 6d) largely mirror the trends for their functional gradient counterparts. However, the maximum range of the SAaligned structural gradient occurs later at 39.3 years (95% CI 14.8–63.8 years), suggesting that microsctructural differentiation along the SA axis plays out well into adulthood. The maximum gradient range of the VS-aligned structural gradient occurs at 3.4 years (95% CI 2.3–4.5 years), indicating that maximum microstructural differentiation with respect to unimodal processing domain is established during Early life. The maximum range of the MR-aligned structural gradient is achieved at 13.65 years (95% CI 9.5–17.8 years). Overall, structural gradient dispersion peaks earlier than functional gradient dispersion, reaching its maximum at 7.75 years (95% CI 6.5–7.0 years). We also studied structure-function gradient similarity and range at the network level (Figure S4), revealing varied timing across networks and gradients for both range and similarity.

### Gradients are stable with respect to sex

We sought to identify sex differences in global gradient characteristics by examining sex-specific lifespan trajectories of gradient metrics (Figure S5a,c). To do so, we first fit a GAMM to each metric as a function of age for the entire population, and obtained residuals from this population-level fit. We then fit a smooth interaction term (sex-by-age) to these residuals, effectively estimating how male and female trajectories might deviate from the mean curve. Sex-specific metric trajectories were reconstructed by adding these sex-specific residual fits back onto the population mean. To formally assess the presence of sex differences, we examined the p-values associated with each sex-by-age interaction term in the residual model. Neither the male nor the female interaction terms reached significance for gradient dispersion or gradient range in any of our axes of interest. This result indicates insufficient evidence to claim a systematic divergence in the shape or amplitude of the lifespan trajectories between males and females. Additionally, we computed spatial maps comparing the lifespan mean gradients of males and females (Figure S5b). Notably, females showed higher values in the association pole and lower values in the somatosensory pole of the SA axis, indicating more connectivity differentiation along the SA axis in females compared to males.

## DISCUSSION

In this work, we explored how fundamental organizing axes of the functional connectome develop throughout the human lifespan. Our findings offer abundant novel observations concerning the unfurling and refinement of fundamental axes of the functional organization of the cortex and unify decades of developmental FC research under one comprehensive normative chart of gradient organization. Our work recapitulates previous findings concerning the timing and topographical organization of FC and synthesizes these results under the gradient mapping analysis framework. Integrating the vast and diverse body of fMRI literature into an easily interpreted and succinct framework represents a central obstacle to neuroscience; the present study offers a substantial step forward towards achieving this goal. Finally, the charting of normative developmental trajectories is a critical tenant of modern medicine, allowing for the identification of disease onset and the targeting of early treatment. While cortical morphology has been mapped in detail throughout the human lifespan^60^, such charts for the development of FC from days after birth into late old age were lacking until now.

Specifically, we studied the sensory-association (SA), visual-somatosensory (VS), and modulation-representation (MR) gradients of functional connectivity, and found that their developmental trajectories are distinct and nonlinear. We first sought to characterize topographic changes in these axes over the lifespan, crucially observing that gradient topography is largely established by 4 years of age. In keeping with previous observations that infant FC architecture is distinct from that of adults^27, 37^, we observe significantly altered SA and MR gradient topography during early life, indicating weakly developed high-order information processing hierarchies and low levels of the long-range connections that support them. Conversely, the VS axis is fully established during middle childhood and its topography remains largely stable while its global range steadily declines throughout adolescence and adulthood. This reduction in range indicates that the differentiation between the two primary sensory modalities becomes a less salient feature of connectivity organization throughout adulthood.

The SA and MR gradients both enumerate information processing hierarchies crucial to complex cognition, with high-order association and control regions positioned at their apexes, respectively. The shape of the GAMM-fitted gradient embedding when viewed in the SA-MR plane (Figure 2b, bottom) sheds light on the interaction between these two gradients, indicating that regions residing near the modulation pole of the MR axis are highly stratified with respect to the SA axis. This shape is striking, and is distinct from the shape of the SA-VS embedding (Figure 2b, top), underscoring the differing roles of the SA and MR hierarchies. We observe similar nonlinear development in gradient range, dispersion, and topographic similarity to the template gradients for the SA and MR axes, indicating that the lifespan development of these transmodal hierarchical axes of organization are interrelated. Despite qualitative similarity in their global development, the decline in range of the SA axis is more dramatic after mid adulthood when compared with the MR axis. This difference suggests that the relative importance of the MR axis as an organizing motif increases significantly during adulthood and aging.

The VS axis undergoes distinct development compared to SA and MR axes, undergoing rapid expansion during infancy and early childhood, followed by contraction throughout the remainder of the lifespan. The range of the VS axis represents the level of differentiation between primary unimodal processing systems; its increase during infancy and childhood reflects functional differentiation of these systems, while its decrease thereafter indicates a continual reduction in the importance of this axis as an organizing motif of the functional connectome. Our observation that the gradient range of the VS axis decreases throughout late childhood and adolescence is consistent with previous work^17, 61^; however our results differ from these studies in that we observe reordering between the SA and VS axes at 3 years old, far earlier than the switching during early adolescence previously observed^61^. This difference could be attributable to the fact that our study utilized GAMM fitting after group-alignment across the entire lifespan, seeking to characterize topographic and global changes in each axis of interest rather than age-bin based analysis. In contrast to the SA and MR axes, the cortical locations occupying the poles of the VS axis (visual and somatosensory cortex) are spatially contiguous. The decrease in the range of the VS axis in tandem with the increase in the range of SA and MR axes after late childhood therefore reflects a transition from a locally-to a globally-organized functional connectivity architecture, which is consistent with prior studies^27^.

Our analysis of global gradient metrics across the lifespan also yielded several notable results that provide insight into the development of global hierarchical FC architecture. A monotonic increase in dispersion between birth and 13.8 years (Figure 3b) indicates that global connectivity differentiation with respect to the SA, VS, and MR axes expands significantly during this period. Declining gradient range of the VS axis during this same period (Figure 3a), however, underscores that the increasing dispersion during this period is driven primarily by expanding transmodal SA and MR axes. It is also notable that increases in gradient dispersion are modest between birth and 13.8 years, a developmental stage well-known for rapid differentiation of cortical FC^23, 62^. In tandem, expanding gradient range during this period in combination with relatively stable dispersion indicates that the extreme values along each gradient are increasing during this period while not driving a significant change in the average spread of vertices in the embedding space. This trend is potentially explained by increased prominence of transmodal hubs at the association and modulation poles of the SA and MR axes in combination with increasing integration among intermediary and specialized cortical locations. Migration of hubs towards more extreme positions in the embedding space engenders increasing gradient range, while increased integration among intermediary nodes constrains the mean global spread (dispersion) of cortical regions. We also observe monotonic decline in global gradient dispersion after 13.8 years, indicating decreasing FC differentiation during aging. This is consistent with previous observations that FC segregation decreases during later adulthood, while within-system FC and between-system FC decrease and increase, respectively^49, 53^.

Many previous studies have studied FC organization via graph-theoretic metrics^63–65^, while little effort has been made to establish connection between FC gradients and underlying graph architecture. Our analysis sought to partially address this gap by studying the relationship between graph-theoretic metrics, including characteristic path length, and metrics derived from diffusion embedding (DE). We found a particularly interesting relationship between gradient dispersion and characteristic path length, observing a moderate negative correlation (*ρ* = 0.25, *p <* 10^−58^) across subjects (Figure 3d). Dispersion is a measure of global connectivity differentiation independent of any particular gradient axis. Characteristic path length quantifies the average shortest path between vertices in a graph, providing a measure of interconnectedness. We found that increasing characteristic path length tends to correspond to increasing dispersion, indicating that FC graphs with more globally differentiated FC are less interconnected on a global scale. Further, we observe that during infancy, dispersion and characteristic path length tend to be low, indicating that infant FC networks have low-levels of global differentiation and have efficient information transfer. This is potentially explained by the high gradient range of the VS axis during this period in combination with a rapidly developing but weakly expressed SA and MR axis. We also observe that adolescents tend to have the highest characteristic path lengths but in general exhibit a broad range of dispersion values, suggesting that increasing network segregation during adolescence is not always accompanied by commensurate increase in global FC differentiation. During young adulthood, characteristic path lengths are low, while dispersion values are high. Together, these trends indicate that mature FC networks are characterized by highly efficient global information transfer in combination with highly differentiated FC. We observe that mean functional connectivity degree increases significantly during infancy and early childhood in tandem with the topographic establishment of our three gradient axes. The decline in FC during adolescence coincides with the rapid reduction in the gradient range of the VS axis during this period, suggesting that global functional connectivity strength is concentrated in unimodal cortex throughout development. Interestingly, the decline in global connectivity strength occurs well before maximum differentiation along the SA and MR axes are achieved, indicating that the growth of these axes is not driven primarily by globally increasing functional connectivity strength.

We also sought to characterize how gradient organization evolves across the human lifespan with respect to resting-state networks (RSNs). We analyzed the trajectories of networks from the well-studied 7-network parcellation of Yeo et al.^57^, characterizing the trajectories and distributions of their constituent vertices in our gradient space. These analyses revealed an intricate and complex sequencing of network development with respect to the SA, VS, and MR axes, with differential trajectories across networks and across gradient axes. Network trajectories along the SA and VS axis are simple and understandable. Motion of each network along these axes coincides with changes in SA and VS range, driving expansion and contraction of the separation of networks at either pole. Importantly, there is very little reordering of the position of networks along the SA and VS axes, indicating a relatively low degree of topographic reorganization with respect to these gradients. In contrast, network centroids along the MR axis undergo significant reordering, with differential timing for maturation across networks. This complexity suggests that the MR axis demands significant topographic reorganization of FC throughout development, reaching maturity at the latest stage compared to SA and VS axes. Interestingly, dorsal and ventral attention networks (DAN and VAN) vary significantly in their developmental timing on the MR axis, with the DAN reaching its maximum MR value during late childhood, and the VAN attaining its maximum value during middle adulthood. This suggests that during early life, refinement of the top-down attention mechanisms associated with the DAN takes center stage, with the development of bottom-up attention (VAN) systems playing out over a more protracted period. Further research on the differential development of these two attention systems and their interrelationships may provide further insight to the cognitive significance of these findings.

Several networks undergo opposite developmental trends in mean value on the MR axis. Namely, the DAN and DMN rapidly separate during infancy and childhood, and move to a more central position throughout the remainder of the lifespan. Similarly, the VAN and somatomotor networks are proximal during infancy, and undergo a prolonged period of increasing MR separation through mid adulthood. This is particularly striking, as it indicates that the increasing differentiation between modulation and representation loci is actualized via increasing contrast between the spatially adjacent premotor and ventral attention networks. Further, the prolonged maturation of the VAN indicates that the development of bottom-up attention and its relationship with primary motor systems continues to mature throughout adulthood, scaffolding a crucial information processing hierarchy.

Our analysis of gradients of cortical morphology and microstructure revealed distinct trajectories for structure-function alignment across the SA, VS, and MR axes. Interestingly, through our gradient-based analysis, we observe that structure-function coupling decreases with respect to all three axes across the lifespan. This reduction is the most dramatic for the SA and MR axes, representing high-order information processing hierarchies. In contrast, the decrease in similarity between the VS axis and its structural counterpart is less dramatic across the lifespan. One potential explanation for this is that microstructural differentiation between visual and somatosensory cortex is pronounced throughout the lifespan. Further, we hypothesize that the high structure-function similarity values for SA and MR axes during infancy could be the result of underdeveloped functional gradients which are initially underpinned by patterns of variation in microstructure which are present at birth. Lifelong decrease in structure-function tethering of the SA and MR gradients indicates that these axes are increasingly driven by global patterns of functional coactivation that do not have a strong basis in microstructural differentiation.

There is strong evidence that gradients of FC, microstructure, and gene expression represent crucial organizing axes of the cerebral cortex^2^. Still, recent pushback about the power of gradients as a comprehensive framework for understanding topographic organization of the cortex bears mentioning. The principle of arealization, whereby the cortex is divided into discrete areas which are differentiated by cytoarchitecture and connectivity, is in some ways at odds with the claim that gradients represent fundamental organizing axes on an individual level^66^. Gradients of functional connectivity typically do not smoothly vary with respect to space on an individual level, reflecting instead sharp boundaries between functional systems. At a population level, however, gradients are smooth and excel at describing functional topography. The interpretation of FC gradients in individuals remains an important area of future research, however the present study focuses on large-scale population-level effects.

### Limitations, outstanding questions, and future directions

It is important to note that there are many degrees of freedom in our methodology, as the present work is fundamentally exploratory. The use of a template gradient set and the subsequent alignment of individual subject gradients to that template prior to GAMM fitting was crucial to standardizing gradient architecture across the lifespan so that the same characteristics of connectivity could be examined at all ages. However, this implies that there exist global organizational features of connectivity that persist from birth to death. The computation of a global lifespan template using weighted principal component analysis applied to all subject gradients is inherently biased towards periods of stable FC gradients. The alignment of infant and childhood gradient sets to this lifespan template yields the best representation of the adult connectivity features present in each infant or child, irrespective of whether those features are principal organizing connectivity motifs. In light of this, our analysis reveals how FC in early life converges towards adult FC with respect to SA, VS, and MR axes, rather than exploring how organizing motifs of FC themselves evolve during early development. Although subtle, this distinction is worth considering and its implications are worth exploring in future studies.

## METHODS

### Materials

Our subjects were selected from 5 imaging datasets: the Baby Connectome Project (BCP)^67^, the development, young-adult, and aging cohorts of the Human Connectome Project (HCP-D, HCP-YA, HCP-A)^68^, and the Healthy Brain Network (HBN) dataset^69^. Subjects in the BCP ranged from 16 days to 6 years old, with 557 subjects scanned at 1,095 unique timepoints prior to gradient-based quality control. We used 652 subjects (age 5.58–21.92 years) from the HCP-D, 1,206 subjects (age 22–37 years) from the HCP-YA, 725 subjects (age 36–100 years) from the HCP-A, and 771 subjects from the HBN (age 5.6–21.9 years). Across all four cohorts, our dataset included 3,951 subjects.

### Quality Control

We excluded infant data with excessive motion, cutting the BCP dataset from 557 subjects with 1,095 timepoints to 343 subjects with 760 timepoints. Motion parameters of all subjects included in our main analysis are displayed in Table S2. For the entire lifespan dataset, we employed visual and clustering-based gradient quality control, excluding problematic gradients from the main analysis. Using this procedure, we excluded 1 additional datapoint from the BCP for a total of 759 timepoints and 343 subjects (158 males, 176 females, 9 unreported). We excluded 2 sets of gradients from the HCP-D for a total of 650 subjects (301 males, 349 females). We retained all data from the HBN dataset (770 subjects, 458 males, 275 females, 37 unreported). We excluded 25 sets of gradients from the HCP-YA for a total of 1,068 subjects (482 male, 586 females). We excluded no data from the HCP-A (725 subjects, 319 males, 406 females). These decisions were justified by examination of subject gradients as well as their associated metrics (dispersion, range, similarity to template) in combination with the clustering procedure described below. In total, our final analysis included 3,972 unique gradient sets from 3,556 subjects. Figure S7 displays the distribution of subject ages by cohort following quality control.

### Data Preprocessing – Functional MRI

We employed an rs-fMRI preprocessing pipeline that is consistent with the HCP^70^. The preprocessing pipeline includes (i) head motion correction using FSL mcflirt; EPI distortion correction using FSL topup to generate distortion correction deformation fields using a pair of reverse phase-encoded FieldMaps (i.e., AP and PA or LR and RL); (iii) rigid (6 DOF) registration of SBref to Fieldmaps; (iv) rigid boundary-based registration (BBR) of distortion corrected SBref image to the corresponding T1-weighted images, with prealignment using a mutual information cost function; and (v) one-step re-sampling, i.e., combining all of the deformation fields and translation matrices above and applying them to the raw 4D rs-fMRI data, resulting in a motion and distortion corrected rs-fMRI sequence in each subject’s native space (T1w space).

We denoised the fMRI data before further analysis. First, we detrended the data via high-pass filtering with a cutoff frequency of 0.001 Hz to remove slow signal drift. Next, we utilized Independent Component Analysis based Automatic Removal Of Motion Artifacts (ICA-AROMA) denoising^71^ to remove any residual motion artifacts. This involved performing a 150-component ICA on the fMRI data and classifying each component as either BOLD signal or artifact based on various criteria such as high-frequency contents, motion parameters, edges, and CSF fractions. Independent components classified as artifacts are (aggressively) removed via regression.

### Data Preprocessing – Structural and Diffusion MRI

Structural data underwent automated quality assessment^72^, inhomogeneity correction, and linear transformation of T2w image to respective T1w image. We then delineated white matter, gray matter, and the cerebrospinal fluid with automated segmentation. Usable diffusion data were selected using a deep-learning based semi-automated QC^73^. Then we performed correction of signal drop-out, pileup, and eddy-current and susceptibility induced distortions^74^. Co-registration between dMRI and sMRI using multi-channel non-linear registration was performed with ANTs. Finally, manual visual inspections were done to ensure quality.

In order to establish a consistent coordinate system across the entire lifespan, we utilized in-house lifespan surface atlases extended from the work of Ahmad et al.^75^ The reconstructed cortical surfaces of each individual were first registered to age-matched cortical surface atlases to get vertex-wise correspondence among all the surfaces across the lifespan. Cortical surfaces were mapped onto unit sphere and Spherical Demons^76^ was used to perform registration between spherical surfaces of the individuals and the age-matched surface atlases. The vertex-wise correspondence established as a result of spherical registration was then propagated to the white and pial surfaces by leveraging one-to-one correspondence between the spherical and standard representation of the white and pial surfaces.

Structural and diffusion indices were computed in the same manner for all cohorts. Specifically, cortical thickness is defined as the Euclidean distance between two correspondent vertices on the white and pial surfaces and myelination is quantified using the ratio between T1w and T2w images^77^. Spherical Mean Spectrum Imaging (SMSI)^78, 79^ was used to calculate intra-cellular volume fraction, extra-cellular volume fraction, intra-soma volume fraction, microscopic fractional anisotropy, microscopic mean diffusivity, microscopic anisotropy index, and orientation coherence index. Diffusion Tensor Imaging (DTI) was used to calculate fractional anisotropy and mean diffusivity.

### Functional Gradients

FC matrices were computed from the vertex-wise fMRI timeseries for each acquistion from each subject using the pairwise Pearson correlation coefficient. We then computed the mean FC matrix for each subject, subjected the mean FC matrix to row-wise thresholding (top 10 percent of connections retained), and computed the normalized angle matrix in keeping with previous gradient literature^1, 18, 34, 80^. We computed 10 FC gradients for each subject using the diffusion embedding implementation in BrainSpace^34^ using the normalized angle matrix as the input affinity matrix. In order to align subject-specific gradient axes consistently across the lifespan, we computed a set of “template gradients” via weighted principal component analysis (WPCA) of all subjects’ gradients. We began by applying a controlled non-linear transformation to each subject’s age, taking the square root of age to capture rapid changes in early life. The transformed ages were partitioned into ten equally spaced bins on the transformed scale and then mapped back to the original age units, yielding narrower bins (and thus finer sampling) for younger subjects. Each bin was assigned an equal total weight, and that weight was uniformly distributed among the subjects within the bin. After normalizing these weights to sum to 1 across all subjects, we constructed a single matrix of dimension (*N*_subj_ × *N*_grad_ × *N*_vert_) by stacking each subject’s set of gradient maps (i.e., *N*_grad_ per subject). We then standardized each vertex column (z-scoring across all subject–gradient observations) before performing WPCA. In WPCA, each row (representing one subject–gradient combination) was multiplied by its subject-specific weight, ensuring that less densely sampled or younger age bins had comparable overall influence relative to bins with many subjects. We extracted the top principal axes of variation (the first ten), and used the first three for subsequent analyses. This weighting strategy mitigates sampling imbalances and preserves important developmental transitions, ensuring that the template gradients capture large-scale connectivity variation equitably across the entire lifespan.

Next, we aligned the gradient sets of all subjects to this template embedding space using one-shot Procrustes alignment^34^ without scaling or mean centering, after which gradients of the same order were correspondent across all subjects. For quality control, we visually inspected individual subject gradients and identified gradients that were dominated by noise and appeared biologically meaningless. We further applied K-means clustering to the subject gradients, and identified a cluster of 25 subjects with corrupted gradients. In total, we excluded the gradients of 28 subjects, leaving a total of 3,972 sets of individual subject gradients.

### Structural Gradients

We also computed cortical gradients based on microstructural features to assess structure-function coupling across the lifespan by applying diffusion embedding to morphometric similarity networks (MSN)^81^. Our structural features included 11 measures: cortical thickness, myelination, intra-cellular volume fraction, extra-cellular volume fraction, intra-soma volume fraction, fractional anisotropy, mean diffusivity, microscopic fractional anisotropy, microscopic mean diffusivity, microscopic anisotropy index, and orientation coherence index For each subject, we constructed feature vectors containing these features at each cortical surface vertex, and computed a structural affinity matrix using pairwise normalized angle between structural feature vectors. We then used this affinity matrix as input to the diffusion map embedding algorithm to obtain structural gradients. In order to compare structural and functional gradients across the lifespan, we aligned each subject’s structural gradients to that subject’s functional gradients, which were previously aligned to the functional template gradients comprised of the SA, VS, and MR axes. All subsequent comparisons between structural and functional gradients were based on these aligned structural gradients.

### Gradient Measures

To fully characterize how gradient architecture evolves throughout the human lifespan, we employ and define a number of quantitative measures of their properties. The Euclidean distance between vertices in a diffusion embedding approximates the geodesic distance in the underlying graph from which it was computed. Thus, the Euclidean distance between vertices in our gradient embedding space approximates connectivity differentiation between those vertices. To measure the global degree of FC differentiation in a subject’s FC diffusion embedding, we define gradient dispersion as the mean Euclidean distance between the embedding coordinate of all vertices and the centroid of the embedding. For a diffusion embedding with embedding coordinates *ξ*_*i*_ = (*x*_*i*_, *y*_*i*_, *z*_*i*_) for each vertex *i* where *x, y*, and *z* refer to the SA, VS, and MR gradient value, respectively, we define dispersion among *N*_vert_ vertices as 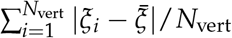. Within-network dispersion was computed analogously for constituent vertices of each of the 7 RSNs^57^.

To characterize the degree of FC differentiation along each gradient axis individually, we compute gradient range. We compute gradient range for each subject and each gradient using the inter-vigintile range (5th percentile to 95th percentile). By using robust statistical measures, we reduce the influence of noise in FC gradients on these global estimates. Gradient range quantifies the degree to which FC is differentiated along a particular gradient axis. Importantly, gradient range is highly correlated with gradient variance (median absolute deviation of gradient value) across subjects (*ρ* = 0.91, 0.86, 0.89 for SA, VS, and MR axes, respectively), indicating consistent range in the diffusion embedding axes prior to scaling by eigenvalue.

The topographic alignment between individual subject gradients and our template gradient was of high interest, allowing us to assess both the quality of the Procrustes alignment procedure and the lifespan changes in gradient topography. To assess this similarity, we compute the cosine similarity between individual subject gradients and the corresponding group-level template gradient.

We also analyzed between-network interactions in our gradient embedding space to probe the lifespan trajectories of inter-network coupling. To this end, we devised a principled measure of network-network embedding distance to quantify the degree of FC similarity between different RSNs. Specifically, between-network embedding distances shown in Figure 5 were computed as the mean Euclidean distance between all possible vertex pairs from two networks normalized by gradient dispersion. Network pairs that are highly segregated in the embedding space correspond to large between-network distances, whereas pairs that are integrated or share similar connectivity profiles have low between-network distances.

### Lifespan Trajectories

Brain development throughout the lifespan is nonlinear. Further, portions of our dataset utilize longitudinal staggered cohort study design. To robustly estimate lifespan trajectories of cortical gradients and gradient-derived metrics, we employed generalized additive mixed models (GAMMs)^35^. This framework is well-suited for neurodevelopmental data as it can flexibly capture nonlinear relationships and incorporate hierarchical random effects, providing stable and biologically informed curve fitting across the wide age range used in this study.

We carried out GAMM fitting and lifespan harmonization of gradient values at the vertex level in order to unify our analysis and streamline downstream computation and modeling of gradient-derived metrics. For each cortical surface vertex, we modeled the gradient value as a smooth function of age, implemented using penalized cubic regression splines and restricted maximum likelihood (REML) for smoothing parameter estimation. Biological changes are more rapid early in life; in order to cater to this, the age variable was transformed by raising age in years to a fractional power, *α*=0.5, improving model stability and distributional assumptions. Subject ID and study cohort were included as random effects to account for repeated measures within individuals in the BCP data and to control for between-cohort differences in data acquisition or processing.

To determine the optimal spline complexity of our GAMMs, we evaluated models with a range of *k*-values. A relatively low *k* (*k* = 4) minimized overfitting and provided stable fits with the majority of cortical surface vertices. We confirmed fit quiality using adjusted *R*^2^ values and visual inspection of residuals. After fitting initial GAMMs to gradient value with this low complexity, we extracted and removed the random intercepts associated with subject ID and study cohort effects. This procedure harmonizes the mean level in gradient value across cohorts, producing mean-adjusted data that reduces site or cohort related biases while preserving biologically meaningful variation.

To address any non-uniformity in the age and cohort distributions of our dataset, we applied a density-based weighting strategy before fitting our final GAMMs to the vertex-wise gradient values. We divided the full transformed age range into equal sized bins and assigned weights to each data point inversely proportional to the bin-specific sample size, ensuring that each age bin contributes equally to our regression and mitigating temporal inhomogeneity. We further scaled these weights by the reciprocal of the cohort sample size to avoid undue influence of large cohorts.

To ensure that our fitted trajectories for gradient values reflect not only the mean trend but also stable variance structure, we controlled for heteroscedasticity using a two-step procedure. First, after mean harmonization using the cohort random effect intercepts of the low-*k* GAMM, we computed vertex-wise residuals and their squares. A second GAMM with low basis complexity and cohort random effects was fit to these squared residuals. The prediction of this variance model estimated how residuals change with age, identifying potential age-dependent heteroscedasticity. For each cohort, scaling the uncorrected residuals to match the predicted variance produced variance-adjusted data, preserving biologically meaningful variability across age while controlling for spurious cohort effects^82^. Finally, we re-fit a GAMM with higher basis complexity (e.g., *k* = 10) to the variance-adjusted data using weights equal to the inverse of the predicted variance at each point, reducing heteroscedasticity in the final model fits.

After harmonization, individual gradient values were mean-corrected for cohort differences and variance-adjusted to address heteroscedasticity. Figure S8 and Figure S9 display diagnostic plots for exemplar vertices, illustrating gradient values and residual distributions before and after the harmonization procedure. Figure S8 exemplifies one of the most significant fits, while Figure S9 exemplifies one of the least significant fits. Figure S11 displays surface maps of effective degrees of freedom values for the final weighted fits for all three gradients. After gradient values were effectively harmonized, analysis of subsequent gradient-derived metrics required no further harmonization. For metrics including gradient dispersion, gradient range, and several networkparcellation-based gradient metrics, GAMMs with no random effects with basis complexities determined by optimal adjusted *R*^2^ and AIC values were fitted with age treated as a smooth term.

To account for the potential confound introduced by arousal state in the BCP data, we divided participants into two cohorts for our GAMM analysis. Specifically, all children aged 3 years or younger were classified as the “sleep cohort” because they were scanned during natural sleep. Participants older than 3 years formed the “wake cohort,” given that they were scanned either while asleep or during a passive movie-watching paradigm^67^. By stratifying the sample in this manner, we aimed to minimize any systematic bias arising from differences in arousal state across these age ranges.

To evaluate the presence of nonlinear age effects at each vertex, we compared a linear model to a generalized additive model (GAM) for each vertex using the harmonized gradient values. First, for a given vertex v, we fit a linear model of the form 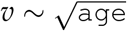, reflecting the hypothesized monotonic but potentially non-uniform relationship between age and the response. Next, we fit a GAM of the form 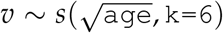, where *s*(·) represents an isotropic smooth function with spline basis dimension k = 6. The GAM was estimated using restricted maximum likelihood (REML) in the mgcv package. Because the linear and GAM formulations are nested (the linear model is a special case of the smoother with effectively 1 degree of freedom), we performed a partial F-test comparing the two models: LM: 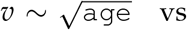. GAM: 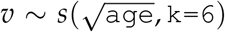. This yields a p-value (*p*_nonlinear_) indicating whether the smoother explains significantly more variance than the linear term alone. We repeated this procedure at each vertex, thus obtaining one *p*_nonlinear_ per vertex. We then applied a false discovery rate (FDR) correction across all vertices’ p-values to control for multiple comparisons. Finally, we defined the proportion of significantly nonlinear vertices as the fraction of vertices for which the FDR-corrected *p*_nonlinear_ was below 0.05. This proportion summarizes the extent to which the age relationship exhibits significant departures from linearity across the cortical surface. We found significantly nonlinear relationships between harmonized gradient value and age for 98% of vertices for the SA axis, 99% of vertices for the VS axis, and 97% of vertices for the MR axis.

We investigated whether males and females exhibited systematically different lifespan trajectories in their global gradient metrics using a two-step GAMM approach. First, to establish a population-level curve, we fit a GAMM to each metric as a function of age (square-root-transformed to account for rapid early-life changes) while ignoring sex. Specifically, 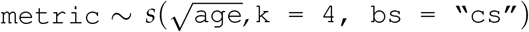, yielding an overall (i.e., sex-agnostic) trajectory. We then computed residuals of each individual’s metric from this population-level fit. Second, to capture sex-specific deviations, we fit a new GAMM to these residuals with a smooth interaction term by sex, 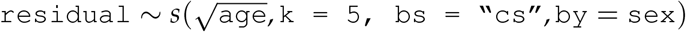, allowing separate smooth functions for males and females. This effectively modeled whether either sex exhibited systematic departures from the population mean trajectory as a function of age. We subsequently obtained sex-specific trajectories by adding each sex’s fitted residual curve back to the population mean fit. We examined the significance of the male- and female-specific smooth terms via their p-values in the GAMM summary. If these terms were significant, it would indicate that one or both sexes diverged from the overall trajectory in a manner that could not be attributed to random variability. Conversely, non-significant terms would suggest insufficient evidence for systematically distinct lifespan trajectories across sexes.

We also investigated whether total brain volume could be a significant confound in lifespan gradient trajectories. To do this, we repeated vertex-wise GAMM fitting for gradient values with total brain volume as a covariate. We then examined the coefficients for the brain volume covariate to assess its impact on the lifespan trajectory of gradient values. Figure S6 shows these coefficients as surface maps, revealing consistently small coefficients across vertices and gradients. Based on this, we conclude that total brain volume is not a significant confound for gradient value across age.

### Estimating Trajectory Extrema

In several cases, it was of interest to obtain estimates of the age at which gradient-derived metrics reached maxima or minima in their lifespan trajectories as an estimate of their maturation age. To obtain uncertainty estimations for the ages at which each metric reached its maximum or minimum value, we used a bootstrapping procedure. We drew 20,000 samples from the posterior distribution of the smooth term’s (age) coefficients, reflecting uncertainty in the model estimates. Each sample represented a possible realization of the age-metric relationship, from which we calculated the fitted metric values across the age sequence. For each bootstrap sample, we identified the age at which the metric achieved its maximum (or minimum) value, thus creating a distribution of ages corresponding to these extreme values across all samples. The 95% confidence intervals (CIs) were then computed from this distribution, providing a probabilistic range within which the true age of the maximum (or minimum) was most likely to be found.

### Mullen Scales of Early Learning

For the BCP participants, we examined the relationships between FC gradient metrics (e.g., dispersion, gradient ranges) and Mullen Scales of Early Learning (MSEL) scores across five subdomains: gross motor, fine motor, visual reception, receptive language, and expressive language. A early learning composite score^67^ was calculated by summing scores from the fine motor, visual reception, receptive language, and expressive language subdomains. Prior to model fitting, participant ages and gradient metrics were z-scored. The Mullen scores were kept in their original scales. For each pair of Mullen score and gradient metric, we fit a linear mixed-effects model using lme4 and lmerTest in R: Mullen Score ~ Age + Gradient Metric + (1 | Subject ID), where subject ID was included as a random intercept to account for repeated measurements. Participants missing data on Mullen scores were excluded on a case-by-case basis. The estimated fixed-effect coefficients (*β*) and the corresponding *p*-values are reported in Table S1.

### Visualization

Figure 1 illustrates our analysis framework and aims to establish visualization conventions to facilitate fast and simple interpretations of our lifespan gradients. First, we associate a color-map with each gradient axis. For the SA, VS, and MR axes, these gradient-specific color-maps span from white to red, green to blue, and black to gray, respectively. To visualize the topography of all three gradients on the cortical surface simultaneously, we devised a color-mapping scheme that combines the gradient-specific color maps such that vertices at the association pole of the SA axis are red, vertices at the modulation pole of the MR axis are gray, and the visual and somatosensory poles of the VS axis are blue and green, respectively. In Figure 1b, we display 3-dimensional embeddings of all three gradients with our 3-dimensional color-map, and establish notation to indicate embedding axis orientation.

Figure 2a utilizes the aforementioned gradient-specific color-maps to chart topographical progression of each gradient in combination with density plots for gradient value. Importantly, we use a constant range with respect to time for the color-maps of surface-based gradient plots. Density plots are computed using kernel density estimation (KDE) applied to the vertex-wise GAMM prediction of gradient value at selected time points. Prior to density estimation, we normalize gradient values across vertices and all timepoints to range between 0 and 1. To showcase temporal changes in both the shape and the scale of density distribution, we use constant horizontal and vertical axis ranges across time.

In Figure 5 (bottom), we display mean between-network embedding distance in matrix form, where mean distance between each network pair is encoded by the color of that cell.

## Supplementary Information

The manuscript contains supplementary material.

## Author Contributions

H. P. Taylor: methodology, software, investigation, visualization, writing - original draft, & writing – review and editing. K. M. Huynh: data curation, methodology, software, & writing – review and editing. K.-H. Thung: data curation, methodology, software, & writing – review and editing. G. Lin: methodology & software. W. Lyu.: resources. W. Lin: resources. S. Ahmad: resources, methodology, software, & writing – review and editing. P.-T. Yap: conceptualization, supervision, funding acquisition, investigation, validation, & writing – review and editing.

## Data Availability

The BCP, HCP-D, HCP-YA, and HCP-A data used in this study are available from the National Institute of Mental Health Data Archive (NDA): https://nda.nih.gov. The HBN data are available from https://healthybrainnetwork.org/.

## Code Availability

Gradient computation and cortical surface visualization were carried out using the BrainSpace toolbox^34^. Volume-to-surface mapping of fMRI timeseries and computation of functional correlation matrices was conducted using Connectome Workbench^83^. Manipulation of FC matrices and computation of gradient-based metrics were carried out in python using standard libraries including Numpy^84^, Scipy^85^, and Sci-Kit Learn^86^. Manipulation of neuroimaging-specific file types was carried out using Nibabel^87^. GAMM fitting was carried out with the MCGV R package^88^, while visualization of lifespan curves and data manipulation in R was carried out using the Tidyverse R libraries^89^.

## Acknowledgements

This work was supported in part by the United States National Institutes of Health (NIH) through grants R01 MH125479, R01 EB008374, and R01 MH133836.

## References

1. Margulies, D. S. et al. Situating the default-mode network along a principal gradient of macroscale cortical organization. Proceedings of the National Academy of Sciences of the United States of America 113, 12574–12579 (2016).

2. Huntenburg, J. M., Bazin, P. L. & Margulies, D. S. Large-scale gradients in human cortical organization. Trends in Cognitive Sciences (2018).

3. Mesulam, M. M. From sensation to cognition. Brain (1998).

4. Buckner, R. L. & Krienen, F. M. The evolution of distributed association networks in the human brain (2013).

5. Sydnor, V. J. et al. Neurodevelopment of the association cortices: Patterns, mechanisms, and implications for psychopathology. Neuron 109, 2820–2846 (2021). URL 10.1016/j.neuron.2021.06.016.

6. Sepulcre, J., Sabuncu, M. R., Yeo, T. B., Liu, H. & Johnson, K. A. Stepwise connectivity of the modal cortex reveals the multimodal organization of the human brain. Journal of Neuroscience (2012).

7. Vázquez-Rodríguez, B. et al. Gradients of structure–function tethering across neocortex. Proceedings of the National Academy of Sciences of the United States of America (2019).

8. Nee, D. E. Integrative frontal-parietal dynamics supporting cognitive control. eLife (2021).

9. Parkes, L., Satterthwaite, T. D. & Bassett, D. S. Towards precise resting-state fmri biomarkers in psychiatry: synthesizing developments in transdiagnostic research, dimensional models of psychopathology, and normative neurodevelopment. Current Opinion in Neurobiology 65, 120–128 (2020).

10. Sporns, O. Graph theory methods: Applications in brain networks. Dialogues in Clinical Neuroscience 20, 111–120 (2018).

11. De Rosa, A. P. et al. Functional gradients reveal cortical hierarchy changes in multiple sclerosis. Human Brain Mapping 45 (2024). URL 10.1002/hbm.26678.

12. Shen, Y. et al. Functional connectivity gradients of the cingulate cortex. Communications Biology 6, 650 (2023). URL 10.1038/s42003-023-05029-0.

13. Smallwood, J. et al. The default mode network in cognition: a topographical perspective. Nature Reviews Neuroscience 22, 503–513 (2021).

14. Zhang, J. et al. Intrinsic functional connectivity is organized as three interdependent gradients. Scientific Reports (2019).

15. Moore, J. W. et al. Gradient organisation of functional connectivity within resting state networks is present from 25 weeks gestation in the human fetal brain. eLife (2023).

16. Park, S., Haak, K., Oldham, S. et al. A shifting role of thalamocortical connectivity in the emergence of cortical functional organization. Nature Neuroscience 27, 1609–1619 (2024). URL 10.1038/s41593-024-01679-3.

17. Xia, Y. et al. Development of functional connectome gradients during childhood and adolescence. Science Bulletin 67, 1049–1061 (2022).

18. Bethlehem, R. A. et al. Dispersion of functional gradients across the adult lifespan. NeuroImage 222, 117299 (2020).

19. Fan, Y. et al. Brain anatomical networks in early human brain development. NeuroImage (2011).

20. Ball, G. et al. Rich-club organization of the newborn human brain. Proceedings of the National Academy of Sciences of the United States of America 111, 7456–7461 (2014).

21. Gao, W. et al. Evidence on the emergence of the brain’s default network from 2-week-old to 2-year-old healthy pediatric subjects. Proceedings of the National Academy of Sciences of the United States of America (2009).

22. Zhang, H., Shen, D. & Lin, W. Resting-state functional MRI studies on infant brains: A decade of gap-filling efforts. NeuroImage 185, 664–684 (2019).

23. Gao, W., Alcauter, S., Smith, J. K., Gilmore, J. H. & Lin, W. Development of human brain cortical network architecture during infancy. Brain Structure and Function (2015).

24. Wang, R. et al. Segregation, integration, and balance of large-scale resting brain networks configure different cognitive abilities. Proceedings of the National Academy of Sciences of the United States of America (2021).

25. Whitaker, K. J. et al. Adolescence is associated with genomically patterned consolidation of the hubs of the human brain connectome. Proceedings of the National Academy of Sciences of the United States of America (2016).

26. Park, B. Y. et al. An expanding manifold in transmodal regions characterizes adolescent reconfiguration of structural connectome organization. eLife (2021).

27. Fair, D. A. et al. Functional brain networks develop from a “local to distributed” organization. PLoS Computational Biology (2009).

28. Luna, B., Padmanabhan, A. & O’Hearn, K. What has fMRI told us about the development of cognitive control through adolescence? Brain and Cognition (2010).

29. Baum, G. L. et al. Modular segregation of structural brain networks supports the development of executive function in youth. Current Biology 27, 1561–1572 (2017).

30. Wendelken, C., Ferrer, E., Whitaker, K. J. & Bunge, S. A. Fronto-parietal network reconfiguration supports the development of reasoning ability. Cerebral Cortex (2016).

31. Geerligs, L., Renken, R. J., Saliasi, E., Maurits, N. M. & Lorist, M. M. A brain-wide study of age-related changes in functional connectivity. Cerebral Cortex (2015).

32. Egimendia, A. et al. Aging reduces the functional brain networks strength—a resting state fMRI study of healthy mouse brain. Frontiers in Aging Neuroscience (2019).

33. Heckner, M. K. et al. The aging brain and executive functions revisited: Implications from meta-analytic and functional-connectivity evidence. Journal of Cognitive Neuroscience (2021).

34. Vos de Wael, R. et al. BrainSpace: a toolbox for the analysis of macroscale gradients in neuroimaging and connectomics datasets. Communications Biology (2020).

35. Chen, C. Generalized additive mixed models. Communications in Statistics - Theory and Methods 29, 1257–1271 (2000).

36. Nazari, R. & Salehi, M. Early development of the functional brain network in newborns. Brain Structure and Function (2023).

37. Yin, W. et al. The emergence of a functionally flexible brain during early infancy. Proceedings of the National Academy of Sciences of the United States of America (2020).

38. Larivière, S. et al. Multiscale structure-function gradients in the neonatal connectome. Cerebral Cortex 30, 47–58 (2020).

39. Schaefer, A. et al. Local-global parcellation of the human cerebral cortex from intrinsic functional connectivity MRI. Cerebral Cortex (2018).

40. Supekar, K., Musen, M. & Menon, V. Development of large-scale functional brain networks in children. PLoS Biology (2009).

41. Keller, A. S. et al. Hierarchical functional system development supports executive function. Trends in Cognitive Sciences 1–15 (2022).

42. Power, J. D. & Petersen, S. E. Control-related systems in the human brain. Current Opinion in Neurobiology (2013).

43. Parlatini, V. et al. Functional segregation and integration within fronto-parietal networks. NeuroImage (2017).

44. Pozuelos, J. P., Paz-Alonso, P. M., Castillo, A., Fuentes, L. J. & Rueda, M. R. Development of attention networks and their interactions in childhood. Developmental Psychology 50, 2405– 2415 (2014).

45. Abrol, A. et al. Developmental and aging resting functional magnetic resonance imaging brain state adaptations in adolescents and adults: A large n (>47k) study. Human Brain Mapping 44, 2158–2175 (2023).

46. Sydnor, V. J. et al. Intrinsic activity develops along a sensorimotor-association cortical axis in youth. Nature Neuroscience 26, 638–649 (2022).

47. Fair, D. A. et al. Development of distinct control networks through segregation and integration. Proceedings of the National Academy of Sciences of the United States of America (2007).

48. Vogel, A. C., Power, J. D., Petersen, S. E. & Schlaggar, B. L. Development of the brain’s functional network architecture. Neuropsychology Review 20, 362–375 (2010).

49. Chan, M. Y., Park, D. C., Savalia, N. K., Petersen, S. E. & Wig, G. S. Decreased segregation of brain systems across the healthy adult lifespan. Proceedings of the National Academy of Sciences of the United States of America (2014).

50. Sun, L. et al. Functional connectome through the human life span. bioRxiv [Preprint] (2024). 2023.09.12.557193.

51. Achard, S. & Bullmore, E. Efficiency and cost of economical brain functional networks. PLoS Computational Biology 3, 0174–0183 (2007).

52. Meunier, D., Achard, S., Morcom, A. & Bullmore, E. Age-related changes in modular organization of human brain functional networks. NeuroImage (2009).

53. Betzel, R. F. et al. Changes in structural and functional connectivity among resting-state networks across the human lifespan. NeuroImage 102, 345–357 (2014).

54. Katsumi, Y. et al. Correspondence of functional connectivity gradients across human isocortex, cerebellum, and hippocampus. Communications Biology 6, 1–13 (2023).

55. Casey, B. J., Tottenham, N., Liston, C. & Durston, S. Imaging the developing brain: What have we learned about cognitive development? Trends in Cognitive Sciences (2005).

56. Menon, V. Developmental pathways to functional brain networks: Emerging principles. Trends in Cognitive Sciences (2013).

57. Thomas Yeo, B. T. et al. The organization of the human cerebral cortex estimated by intrinsic functional connectivity. Journal of Neurophysiology (2011).

58. Huttenlocher, P. R. & Dabholkar, A. S. Regional differences in synaptogenesis in human cerebral cortex. Journal of Comparative Neurology (1997).

59. Petanjek, Z. et al. Extraordinary neoteny of synaptic spines in the human prefrontal cortex. Proceedings of the National Academy of Sciences of the United States of America (2011).

60. Bethlehem, R. A. I. et al. Brain charts for the human lifespan. Nature 604, 525–533 (2022).

61. Dong, H. M., Margulies, D. S., Zuo, X. N. & Holmes, A. J. Shifting gradients of macroscale cortical organization mark the transition from childhood to adolescence. Proceedings of the National Academy of Sciences of the United States of America (2021).

62. Wen, X. et al. First-year development of modules and hubs in infant brain functional networks. NeuroImage (2019).

63. van den Heuvel, M. P. & Sporns, O. Network hubs in the human brain. Trends in Cognitive Sciences 17, 683–696 (2013).

64. Sporns, O. & Betzel, R. F. Modular brain networks. Annual Review of Psychology (2016).

65. Bassett, D. S. & Bullmore, E. T. Small-world brain networks revisited. Neuroscientist 23, 499–516 (2017).

66. Petersen, S. E., Seitzman, B. A., Nelson, S. M., Wig, G. S. & Gordon, E. M. Principles of cortical areas and their implications for neuroimaging. Neuron 112, 2837–2853 (2024).

67. Howell, B. R. et al. The UNC/UMN baby connectome project (BCP): An overview of the study design and protocol development. NeuroImage (2019).

68. Harms, M. P. et al. Extending the human connectome project across ages: Imaging protocols for the lifespan development and aging projects. NeuroImage (2018).

69. O’Connor, D. et al. The healthy brain network serial scanning initiative: A resource for evaluating inter-individual differences and their reliabilities across scan conditions and sessions (2017).

70. Thung, K.-H., Wu, Z., Wang, L., Lin, W. & Yap, P.-T. Analysis of ICA-AROMA motion denoising on fMRI data in infant cohort. In Annual Meeting of the Organization for Human Brain Mapping (OHBM) (2022).

71. Pruim, R. H. et al. ICA-AROMA: A robust ICA-based strategy for removing motion artifacts from fMRI data. NeuroImage 112, 267–277 (2015).

72. Liu, S. et al. Learning MRI artefact removal with unpaired data. Nature Machine Intelligence 3, 60–67 (2021).

73. Liu, S. et al. Multi-stage image quality assessment of diffusion MRI via semi-supervised nonlocal residual networks. In Medical Image Computing and Computer Assisted Intervention (MICCAI), 521–528 (Springer, 2019).

74. Ahmad, S. et al. Fast correction of eddy-current and susceptibility-induced distortions using rotation-invariant contrasts. In Medical Image Computing and Computer-Assisted Intervention (MICCAI), 34–43 (Springer, 2020).

75. Ahmad, S. et al. Multifaceted atlases of the human brain in its infancy. Nature Methods 20, 55–64 (2023).

76. Yeo, B. et al. Spherical demons: Fast diffeomorphic landmark-free surface registration. IEEE Transactions on Medical Imaging 29, 650–668 (2010).

77. Glasser, M. F. & Van Essen, D. C. Mapping human cortical areas in vivo based on myelin content as revealed by T1-and T2-weighted MRI. Journal of Neuroscience 31, 11597–11616 (2011).

78. Huynh, K. M. et al. Probing tissue microarchitecture of the baby brain via spherical mean spectrum imaging. IEEE Transactions on Medical Imaging 39, 3607 – 3618 (2020).

79. Huynh, K. M., Wu, Y., Ahmad, S. & Yap, P.-T. Microstructure fingerprinting for heterogeneously oriented tissue microenvironments. In Medical Image Computing and Computer-Assisted Intervention (MICCAI) (Springer, 2023).

80. Paquola, C. et al. Microstructural and functional gradients are increasingly dissociated in transmodal cortices. PLoS Biology (2019).

81. Seidlitz, J. et al. Morphometric Similarity Networks Detect Microscale Cortical Organization and Predict Inter-Individual Cognitive Variation. Neuron (2018).

82. Pomponio, R. et al. Harmonization of large MRI datasets for the analysis of brain imaging patterns throughout the lifespan. NeuroImage (2020).

83. Marcus, D. S. et al. Informatics and data mining tools and strategies for the human connectome project. Frontiers in Neuroinformatics 5 (2011).

84. Harris, C. R. et al. Array programming with NumPy. Nature 585, 357–362 (2020).

85. Virtanen, P. et al. SciPy 1.0: Fundamental Algorithms for Scientific Computing in Python. Nature Methods 17, 261–272 (2020).

86. Pedregosa, F. et al. Scikit-learn: Machine learning in Python. Journal of Machine Learning Research 12, 2825–2830 (2011).

87. Brett, M. et al. nipy/nibabel: 5.2.1 (2024). URL 10.5281/zenodo.10714563.

88. Wood, S. Generalized Additive Models: An Introduction with R (Chapman and Hall/CRC, 2017), 2 edn.

89. Wickham, H. et al. Welcome to the tidyverse. Journal of Open Source Software 4, 1686 (2019).

