## Supplementary Info for "Functional Hierarchy of the Human Neocortex from Cradle to Grave"

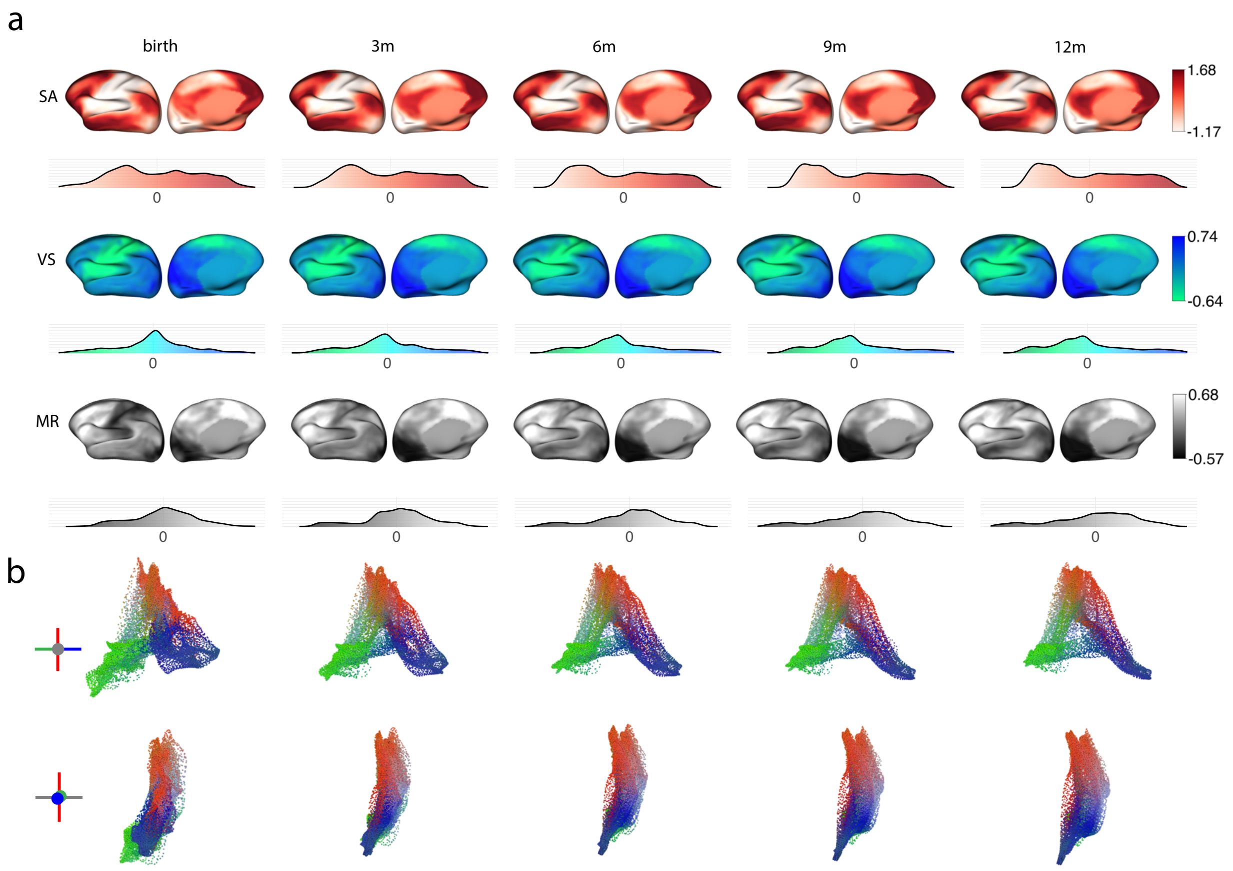


**Fig. S1** | GAMM fitted gradients during the first year of life.

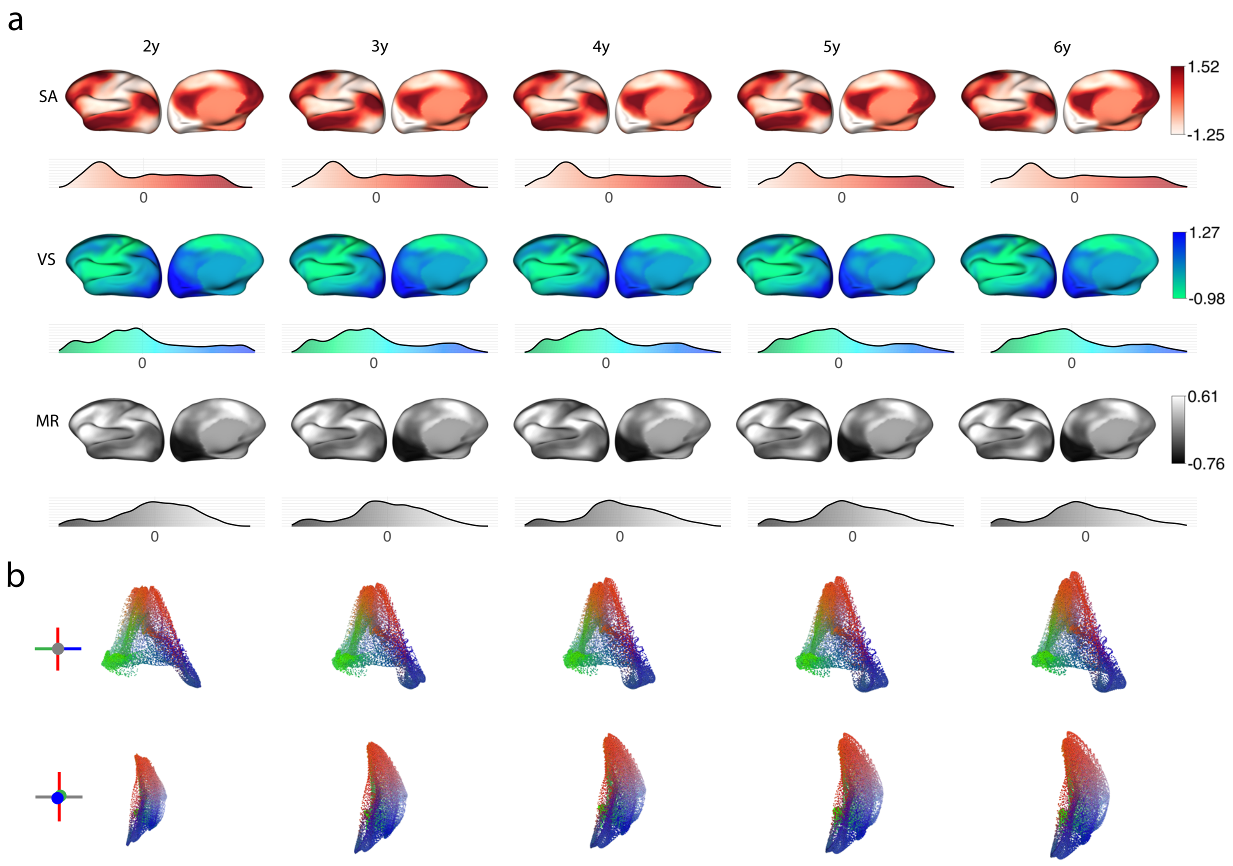


**Fig. S2** | GAMM fitted gradients between 2 and 6 years.

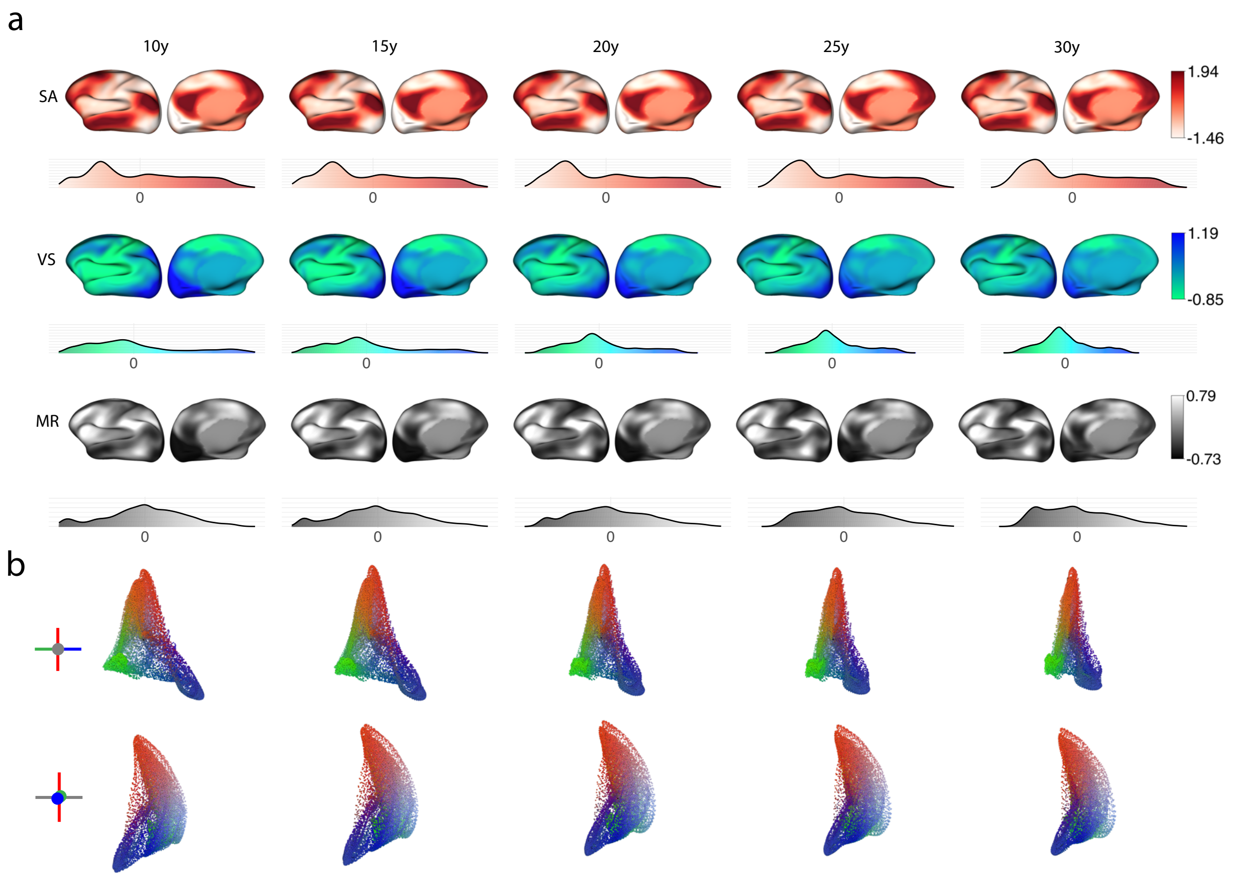


**Fig. S3** | GAMM fitted gradients between 10 and 30 years.

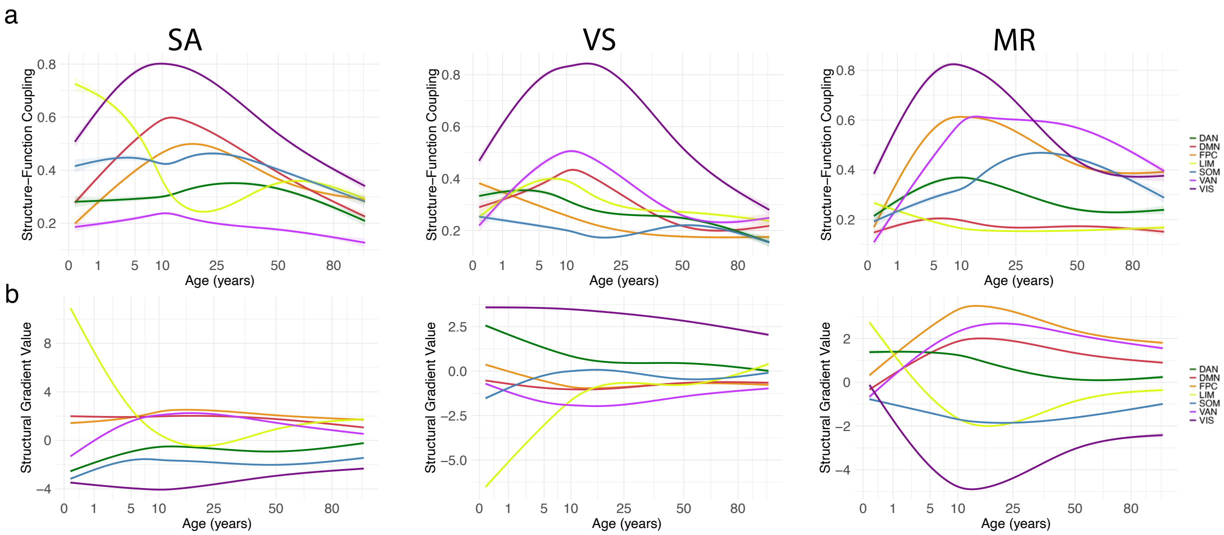


**Fig. S4** | Network-specific **a**, structure-function gradient coupling and **b**, structural gradient values based on the Yeo 7-network parcellation.

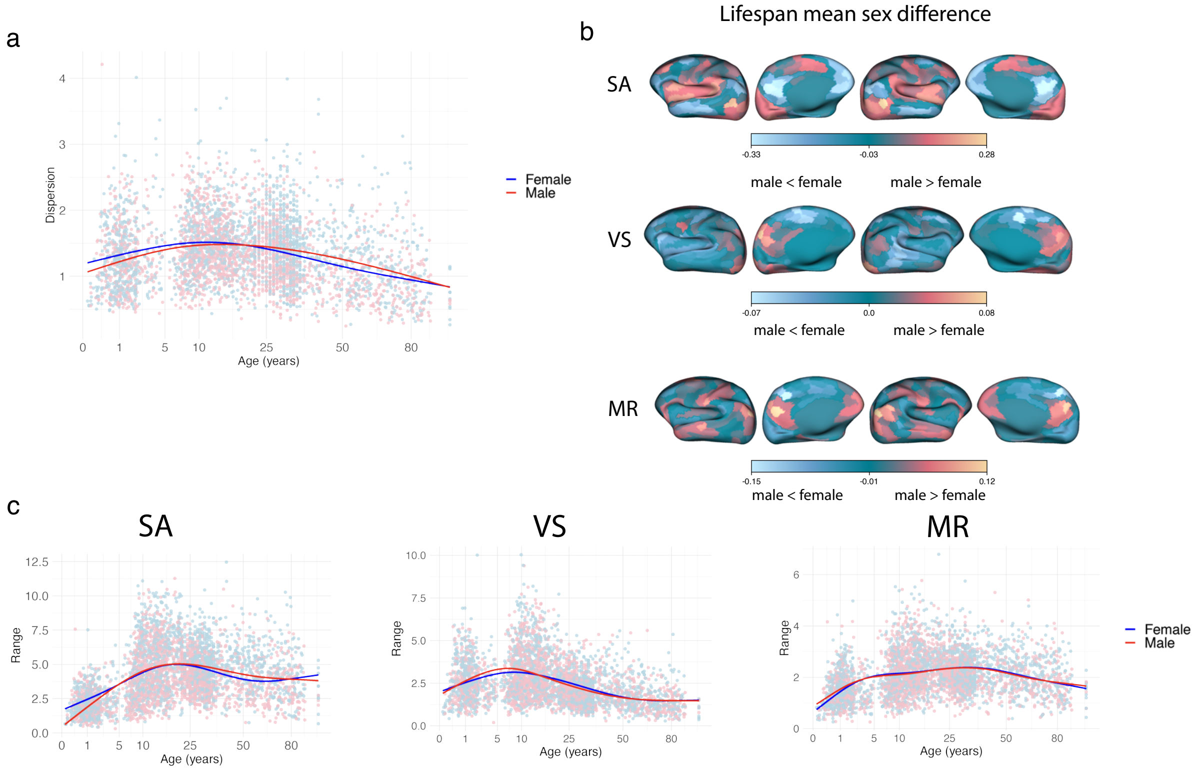


**Fig. S5** | **a**, GAMM fits of gradient dispersion for males and females. **b**, Surface maps of lifespan mean sex difference between parcellated gradient values. **c**, GAMM fits of SA, VS, and MR gradient ranges (from left to right) plotted against age for males and females.

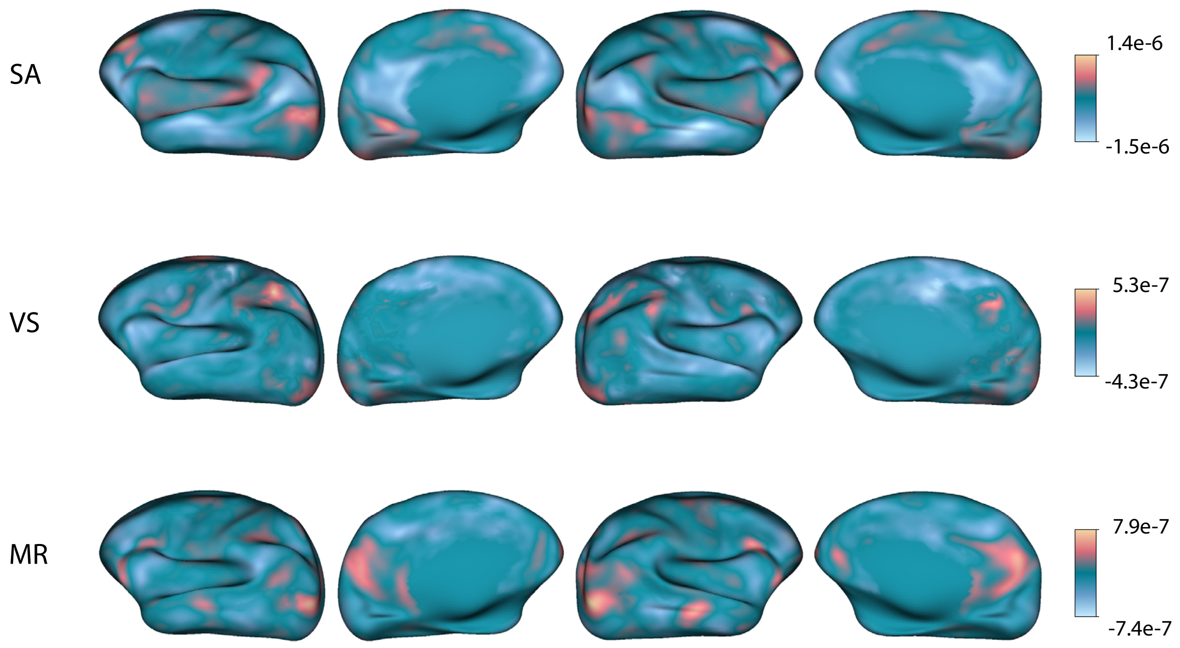


**Fig. S6** | Surface plots showing the effects of the total brain volume on vertex-wise gradient values across age.

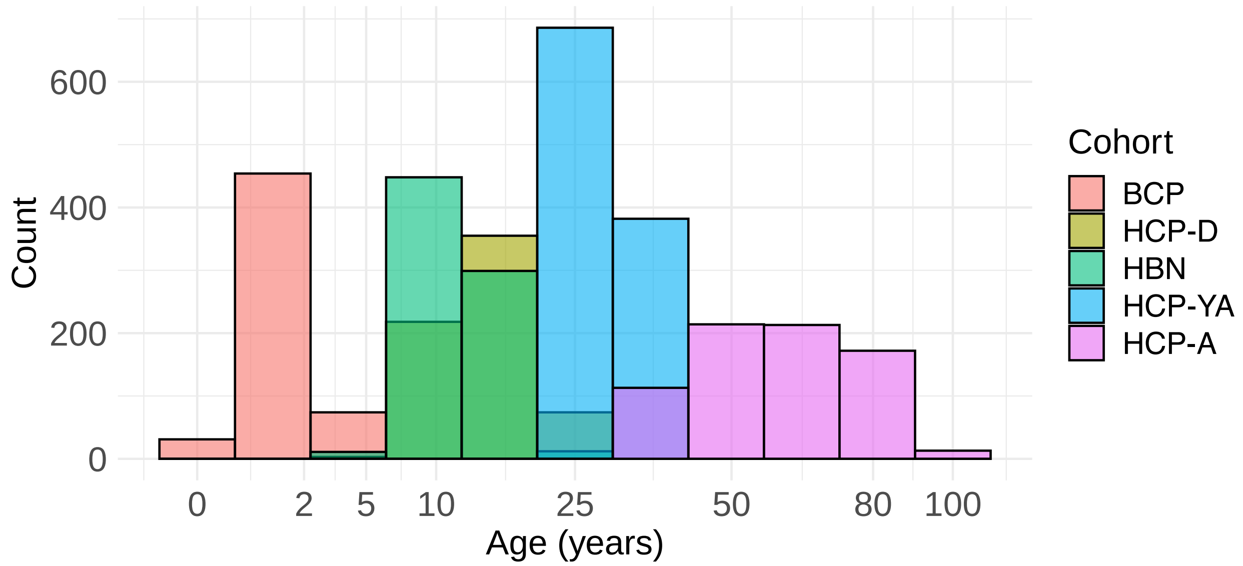


**Fig. S7** | Histogram of subject ages across all cohorts.

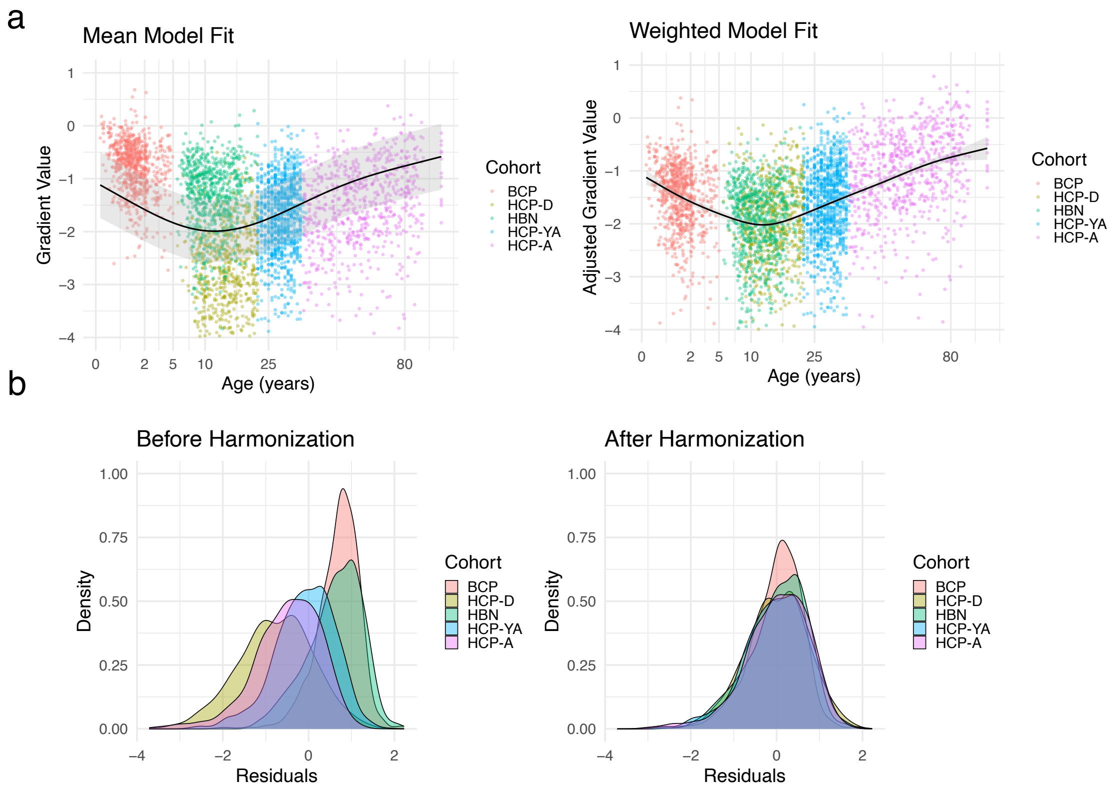


**Fig. S8** | Diagnostic plots for the harmonization procedure for an example vertex with a strong fit. **a**, Gradient values before harmonization for an example vertex with the initial mean model fit (left) and variance and mean-adjusted gradient values with the final weighted model fit (right). **b**, Residual distributions by cohort before (left) and after (right) the harmonization procedure.

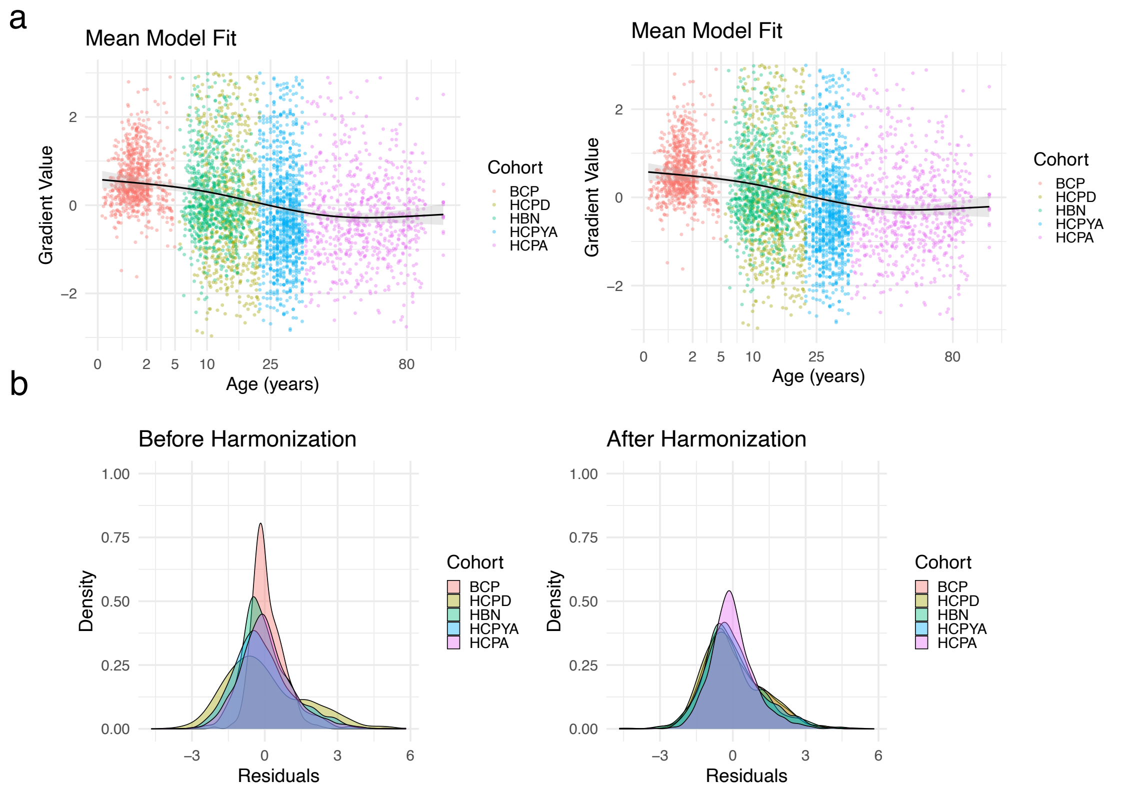


**Fig. S9** | Diagnostic plots for the harmonization procedure for an example vertex with a weaker GAMM fit. **a**, Gradient values before harmonization for an example vertex with the initial mean model fit (left) and variance and mean-adjusted gradient values with the final weighted model fit (right). **b**, Residual distributions by cohort before (left) and after (right) the harmonization procedure.

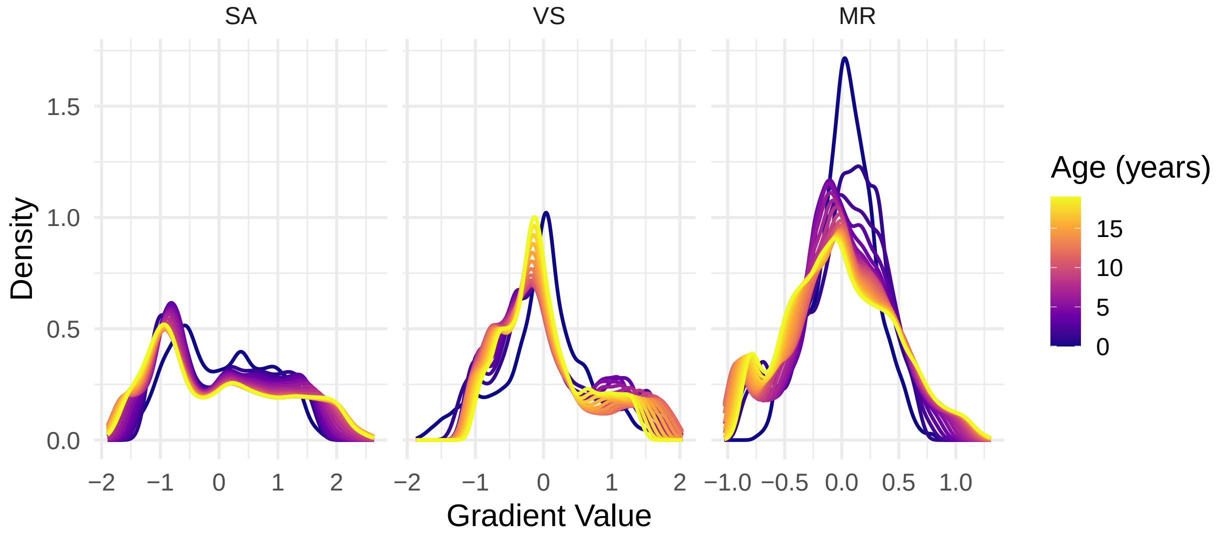


**Fig. S10** | Density plots for gradient value for the first 20 years of life plotted on the same axis and color-mapped to age for SA, VS, and MR gradients.

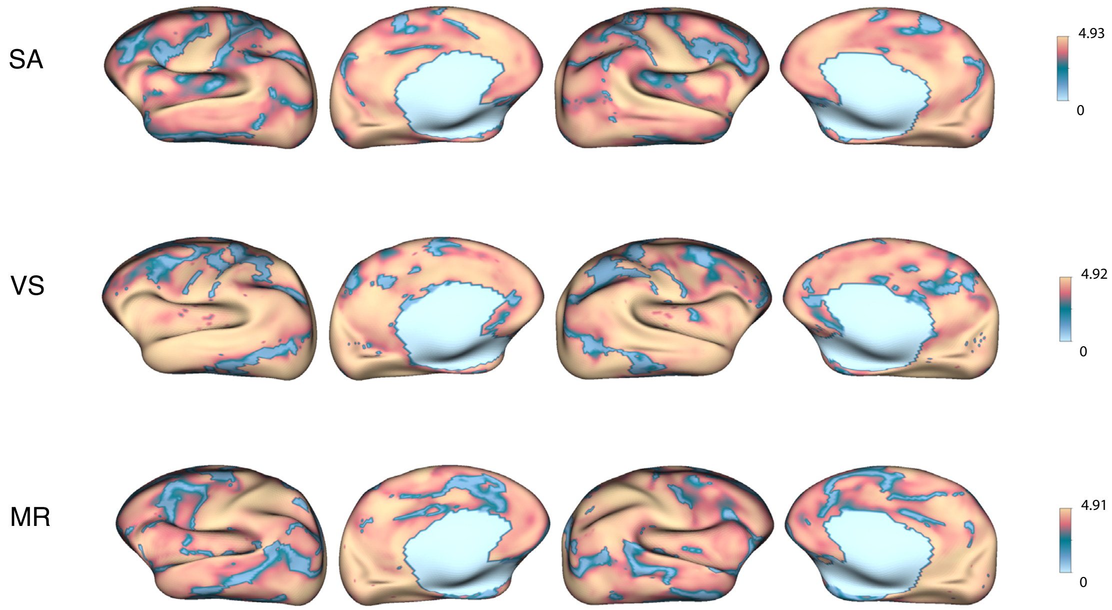


**Fig. S11** | Vertex-wise effective degrees of freedom values for GAMM fits of the SA, VS, and MR gradients.

**Tab. S1** | The relationship between each Mullen score and each gradient metric was examined using linear mixed-effects models, controlling for age and including a random intercept for each participant. Each cell displays the estimated coefficient ($\beta$) for the gradient metric, along with the corresponding p-value.

| Metric | Gross Motor | Fine Motor | Visual | Rec. Language | Exp. Language | Composite |
|---|---|---|---|---|---|---|
| Dispersion | 0.027/0.083 | 0.010/0.517 | 0.008/0.559 | -0.014/0.345 | -0.011/0.437 | -0.001/0.906 |
| SA range | 0.007/0.670 | -0.003/0.856 | 0.000/0.980 | -0.015/0.310 | -0.012/0.402 | -0.007/0.494 |
| VS range | 0.037/0.013 | 0.008/0.614 | 0.017/0.240 | -0.007/0.627 | -0.000/0.980 | 0.005/0.649 |
| MR range | 0.020/0.192 | 0.047/0.002 | 0.017/0.236 | -0.001/0.943 | -0.003/0.853 | 0.016/0.142 |
| SA cossim | 0.007/0.648 | 0.007/0.647 | 0.014/0.322 | -0.002/0.881 | 0.021/0.157 | 0.010/0.339 |
| VS cossim | 0.027/0.075 | 0.027/0.077 | 0.031/0.031 | 0.014/0.354 | 0.010/0.508 | 0.020/0.059 |
| MR cossim | 0.014/0.361 | 0.036/0.021 | 0.028/0.057 | 0.025/0.100 | 0.025/0.101 | 0.027/0.012 |
| Eval 1 | 0.028/0.066 | 0.023/0.119 | 0.020/0.166 | -0.003/0.856 | -0.011/0.429 | 0.007/0.492 |
| Eval 2 | 0.041/0.009 | 0.041/0.007 | 0.017/0.251 | -0.007/0.615 | -0.008/0.577 | 0.010/0.356 |
| Eval 3 | 0.049/0.002 | 0.069/0.000 | 0.031/0.035 | 0.011/0.476 | -0.002/0.879 | 0.027/0.014 |

**Tab. S2** | Cohort-specific QC statistics.

| **Cohort** | **Number of Subjects** | **Number of Scans** | **Median Mean FD** (mm) |
|---|---|---|---|
| BCP | 343 | 1,932 | 0.2684 |
| HCP-D | 650 | 2,505 | 0.1371 |
| HBN | 770 | 1,533 | 0.1804 |
| HCP-YA | 1,068 | 4,121 | 0.1435 |
| HCP-A | 725 | 2,885 | 0.1667 |
